# Dynamical decoration of stabilized-microtubules by tau-proteins

**DOI:** 10.1101/638858

**Authors:** Jordan Hervy, Dominique J. Bicout

## Abstract

Tau is a microtubule-associated protein that regulates axonal transport, stabilizes and spatially organizes microtubules in parallel networks. The Tau-microtubule pair is crucial for maintaining the architecture and integrity of axons. Therefore, it is essential to understand how these two entities interact to ensure and modulate the normal axonal functions. Based on evidence from several published experiments, we have developed a two-dimensional model that describes the interaction between a population of Tau proteins and a stabilized microtubule at the scale of the tubulin dimers (binding sites) as an adsorption-desorption dynamical process in which Tau can bind on the microtubule outer surface via two distinct modes: a longitudinal (along a protofilament) and lateral (across adjacent protofilaments) modes. Such a process yields a dynamical distribution of Tau molecules on the microtubule surface referred to as *microtubule decoration* that we have characterized at the equilibrium using two observables: the total microtubule coverage (density of the Taus on the microtubule surface) and the nearest neighbor distribution of Tau. Using both analytical and numerical approaches, we have derived expressions and computed these observables as a function of key parameters controlling the binding reaction: the stoichiometries of the Taus in the two binding modes, the binding equilibrium constants and the ratio of the Tau concentration to that of microtubule tubulin dimers.

## I. INTRODUCTION

Microtubules are one of the three types of filamentous polymers that constitute the cellular cytoskeleton. A key feature of microtubules is their dynamic nature [1, 2]. This dynamical behaviour, referred to as *dynamic instability*, is exquisitely regulated and is crucial to many cellular activities including cell division, intracellular transport and the establishment and maintenance of cell shape and polarity [3]. Tau (Tubulin Associated Unit) is an important microtubule-regulating protein that is predominantly expressed in axons [4]. This neuronal protein has been reported to cover a large range of fundamental microtubule-related functions. In particular, tau promotes tubulin assembly [5, 6], stabilizes (i.e, regulates) the dynamic instability of microtubules [7, 8], spatially organizes microtubules in a parallel network in axons [9] and can control the axonal transport in regulating the walk of kinesins and dyneins along microtubules [10]. Overall, tau significantly contributes to the stabilization of neuronal microtubules, although the mechanisms underlying these biological functions are still not well understood. Furthermore, appearance of dysfunctions in the couple tau-microtubule has been correlated with numerous neurodegenerative diseases commonly referred as *tauopathies* including Alzheimers, Huntington’s and Pick’s diseases [11–13]. This group of neurodegenerative diseases is characterized by an accumulation of abnormal tau protein in the human brain [14]. Both gain of toxicity and loss of normal function of tau-proteins are though to contribute to the development of *tauopathies* [3, 15].

Because of its important implication in neurodegenerative disorders, tau has been the focus of much study, with a recent emphasis on tau-based therapeutic strategies [16, 17]. To understand how tau ensure the essential normal functions, it is of paramount importance to determine how it interacts with microtubules. In addition to the long-standing experimental effort, simulations of the molecular dynamics of the Tau protein along with a MT section have recently been performed [44]. In the present study, we are interested in the binding reaction between a population of tau-proteins and a stabilized-microtubule. As illustrated in Fig. 1, for a given concentration of tau in the solution and a given concentration of polymerized tubulin dimers forming the microtubule, the reaction yields to a dynamical distribution of tau on the microtubule surface, which we will refer to as *microtubule decoration*. The main objective of this paper is to characterize a such decoration in terms of the mean number of bound-tau and the spatial distribution of tau on the microtubule surface. To study how these two observable quantities vary as functions of key parameters controlling the binding reaction, we have developed a general model based on data from the published literature. To build a reasonable picture of the tau-microtubule interaction that will serve as a basic premise for our model, we considered two aspects. The first is the structure of the microtubule lattice that describes the playground where tau-proteins bind. The second are the characteristics of how tau proteins specifically interact with a stabilized-microtubule that include the tau-protein structure, the location/definition of binding sites (on the microtubule) for tau-proteins, the binding geometry of tau and the binding reaction stoichiometry. A database on what is known on these two aspects was constructed with data from the literature (see supplementary information) and the main results summarized below.

**FIG. 1:**
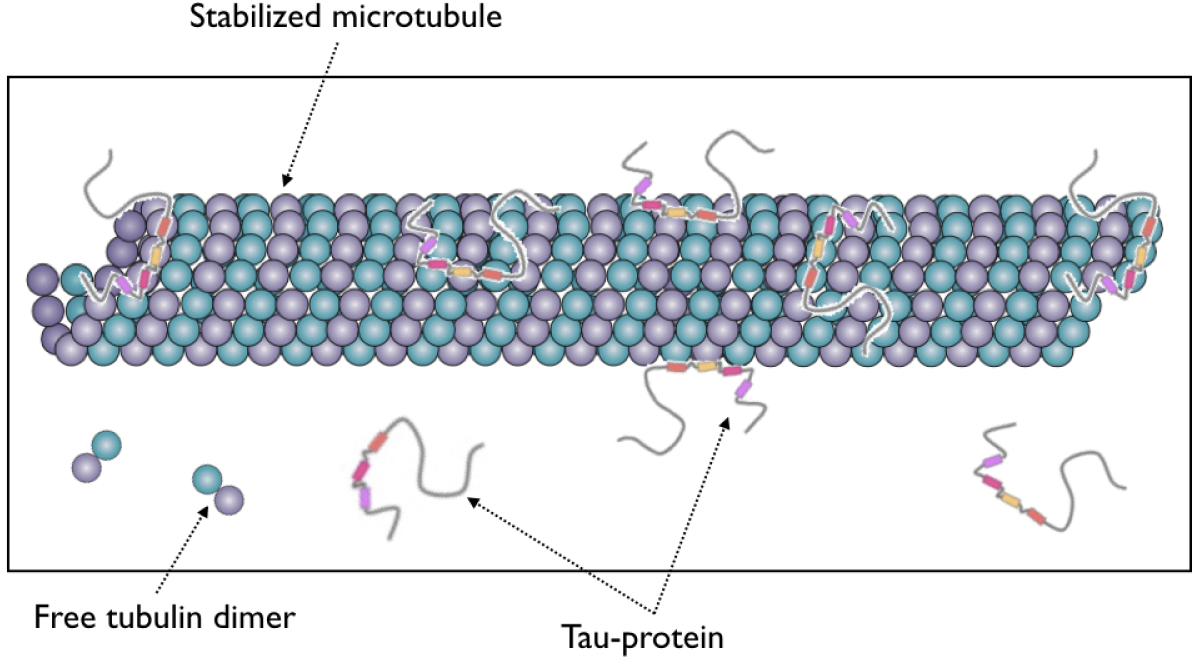
Cartoon showing a population of Tau’s binding on / unbinding from a stabilized microtubule. Adapted from [56] and [57].

## II. LITERATURE ANALYSIS

### Structure of the microtubule lattice

Microtubules (MT) are composed of 8 *nm* long *αβ*-tubulin dimers which are aligned end-to-end to form linear protofilaments [18]. Most of the microtubules assembled *in vitro* and *in vivo* are composed of *p* = 13 protofilaments [19] with a longitudinal shift of 12/13 ≈ 0.92 *nm* between protofilaments, generating a left-handed three-start helix [20–22]. As shown in Fig. 2a,b, the distance separating two protofilaments is about 5 *nm* [20]. At the microscopic level, *αβ*-tubulin heterodimers are packed in a B-type lattice, which has been found to be the most favorable configuration [23]. Interactions between protofilaments in the lattice involve homologous subunits (*α* −*α* and *β* − *β*) except at the seam (i.e., between the first and last protofilament), where a discontinuity exists due to the pitch of three tubulin monomers. As a consequence of such a packing, the microtubule is a polar structure with two extremities, “+ end” exhibiting *β* monomers, and, “-end” exhibits *α* monomers. In this study we assumed that the curvature of the microtubules because of its helical geometry has no effect on the binding of tau molecules and that the 13 protofilaments consisting the microtubule are all identical. Therefore, we use the unfold and flattened bidimensional representation shown in Fig. 2b as an appropriate model of the MT surface for the process of attachment of tau-proteins.

**FIG. 2:**
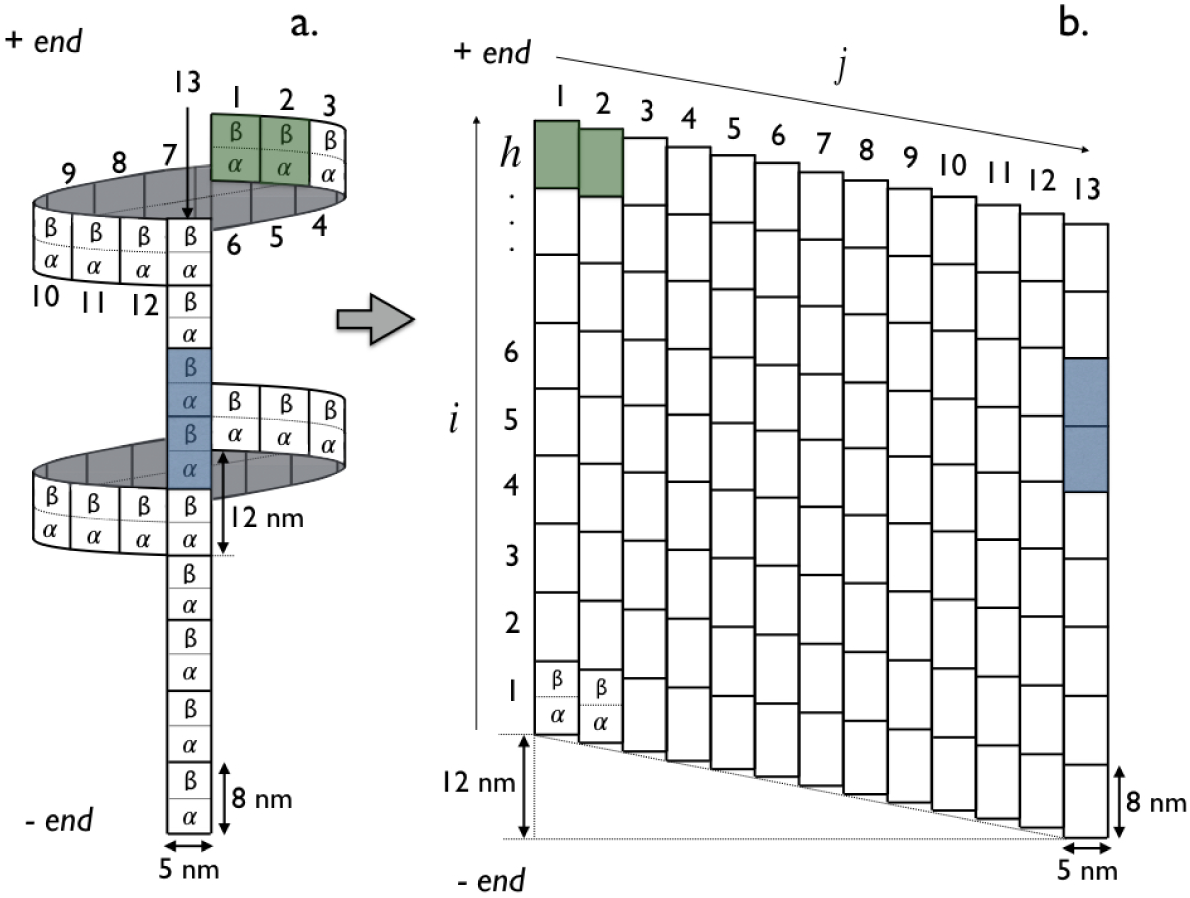
Tau’s binding sites on the microtubule representations (see main text for detailed explanations). “*p*” (blue) and “*h*”(green) modes of binding are illustrated in dark color. (**a**) Three-dimensional representation of the 13− protofilament microtubule. (**b**) Two-dimensional lattice mapping of the 13− protofilament microtubule with *N* = *h*×*p* lattice sites (tubulin dimers) where *h* is the number of *αβ*-tubulin dimers along the protofilament axis *i* and *p* = 13 is the number of protofilaments (or *αβ*-tubulin dimers along the helix axis *j*).

### Interaction tau-microtubule

1. Structure of tau proteins: Tau is a “natively unfolded” molecule with a radius of gyration of 5 − 7 *nm* in solution [18, 24]. There are 6 variants of the tau protein called isoforms which are distinguished by their amino acid sequences. Tau can be regarded as a dipole with two domains of opposite charge, a microtubule-binding domain which includes 3 (tau 3R) or 4 (tau 4R) sequence repeats and a projection domain regulating the spacing between microtubules in axons [25, 26]. It has been shown that repeats bind independently of each other and that the affinity of tau increases with the number of repeats [27].
2. Location of the binding sites: The exact tau-MT binding sites are still not well defined [28]. Comparisons between tau decorated and control microtubules using cryo-electron microscopy revealed binding of tau proteins at the outer microtubule surface [29–31]. This was also supported by atomic force microscopy [32] while a study in 2003 reported a possible binding site at the inner microtubule surface close to the taxol-binding site on *β*-tubulin [33]. Later on, Makrides *et al* [34] suggested that these discrepancies may come from differences in the experimental protocol when adding tau proteins into the solution with either an addition to pre-stabilized MTs or to polymerizing tubulin. However, a recent high resolution cryo-EM study has shown that in both experimental conditions, tau was always bound at the outer MT surface [31]. Moreover, authors in [31] proposed a model in which tau interacts with both *α*- and *β*-tubulin. In this study, we will consider that the binding of tau occurs at the outer MT surface with the *α* − *β*-tubulin dimer as the elementary unit binding site as shown in Fig. 2.
3. Binding modes: The binding mode and the geometry of tau when bound to the MT surface is still very controversial. Some studies [30, 31, 35, 36] have suggested that a tau preferentially adopt an ordered structure aligning along protofilament ridges when bound to the MT, while structures of bound tau crossing adjacent protofilaments were observed as well in [32]. And a combination of high-resolution metal-shadowing and cryo-EM has revealed the existence of longitudinal and lateral bound-tau on the same MT [29]. This latter observation is consistent with a recent study showing that tau promotes the formation of tubulin rings alone and stacks of tubulin rings [37]. In the absence of any further information and to keep generality, we will consider in what follows that a tau-protein is likely to bind on the outer MT surface with two binding modes: a longitudinal mode (“*p*” mode), in which the binding occurs along a single protofilament and, a lateral mode (“*h*” mode), where the binding takes place across adjacent protofilaments along the helix, see Figs. 2 and 3.
4. Stoichiometry: Most of the reported values in the literature seem to converge towards a stoichiometry of 0.5; *ν* = 0.4 [38–40], *ν* = 0.412 [41], *ν* = 0.46 [42] and *ν* = 0.52 [30]. In addition, when the above-mentioned stoichiometries are corrected following the approach in [43], we end up with the value *ν* = 0.5 corresponding to “1” tau molecule for “2” *α* − *β*-tubulin dimers (i.e., 1 tau for 4 tubulin monomers). This experimental evidence is supported by a recent work [44] on the Tau - MT interaction using molecular dynamics simulations that show that bound Tau molecules are in an extended conformation and interact with two *α* − *β*-tubulin dimers on average. However, a recent study suggests that Tau may span up to 4 *α* − *β*-tubulin dimers with thus a smaller stoichiometry of *ν* = 0.25 [31]. Therefore, in this work, we have developed a general model that allows to deal with any stoichiometry. However, the case of *ν* = 1/2 will often be used in illustrations.

To summarize, we will consider from now on that a bound tau molecule, at the outer MT surface, covers 1*/ν* adjacent *α* − *β*-tubulin dimers either along a single protofilament or across adjacent protofilaments. Figure 2 synthetically summarized the premise results for the case of *ν* = 1/2. This picture will be used as a starting point to define the binding rules in the model describing the dynamical decoration of a stabilized-microtubule with a population of tau-proteins.

## III. DECORATION MODEL

To characterize the decoration of microtubules with Tau-proteins, we will be interested in determining the quantities like the coverage of MT by Tau-proteins, the distribution of Tau spacing on the MT surface and the arrangement order of Tau-proteins as functions of key parameters controlling the decoration process. To this end, we describe the binding rules of Tau-proteins on the MT lattice and the kinetic process of attachment-detachment of Tau-proteins.

### Binding rules

We visualize a Tau-molecule as a stem of length *σ* _…_ and extension 0 (both relative to the binding site size) and, thus, denote by *σ*_*p*_ and *σ*_*h*_ (positive integers) lengths or size of a Tau when bound in “*p*” and “*h*” modes, respectively. In this respect, a Tau-molecule bound in “*p*” mode covers (1 + *σ*_*p*_) consecutive binding sites on a single protofilament along the protofilament axis; corresponding to 1 binding site on (1 + *σ*_*p*_) consecutive helices. Likewise, a Tau bound in “*h*” mode covers (1 + *σ*_*h*_) consecutive binding sites on a single helix along the helicoidal axis; corresponding to 1 binding site on (1 + *σ*_*h*_) adjacent protofilaments. The binding of a Tau can only occur on free lattice binding sites; neither partial, nor overlapping, nor stacked bindings are allowed and no “*h*” mode binding at the seam (i.e., crossing protofilaments *j* = 1 and *j* = 13) is allowed. The binding of Tau is a saturable process; accumulation on the MT surface is not possible. The stoichiometry matrix associated with these rules reads as,

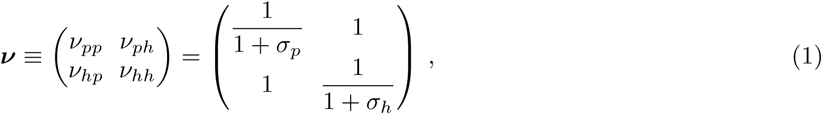

where diagonal elements *ν*_*pp*_ and *ν*_*hh*_ are the stoichiometries of Tau’s bound in “*p*” and “*h*” modes, respectively, and the off-diagonal element *ν*_*ph*_ (*ν*_*hp*_) represents the apparent projection stoichiometry along the helix (protofilament) axis for a Tau bound in “*p*” (“*h*”) mode. These binding rules are illustrated and summarized in Fig. 3 for a microtubule lattice made of *N* = 9 × 13 binding sites in the case of *ν* = 1/2 corresponding to *σ*_*p*_ = *σ*_*h*_ = 1.

**FIG. 3:**
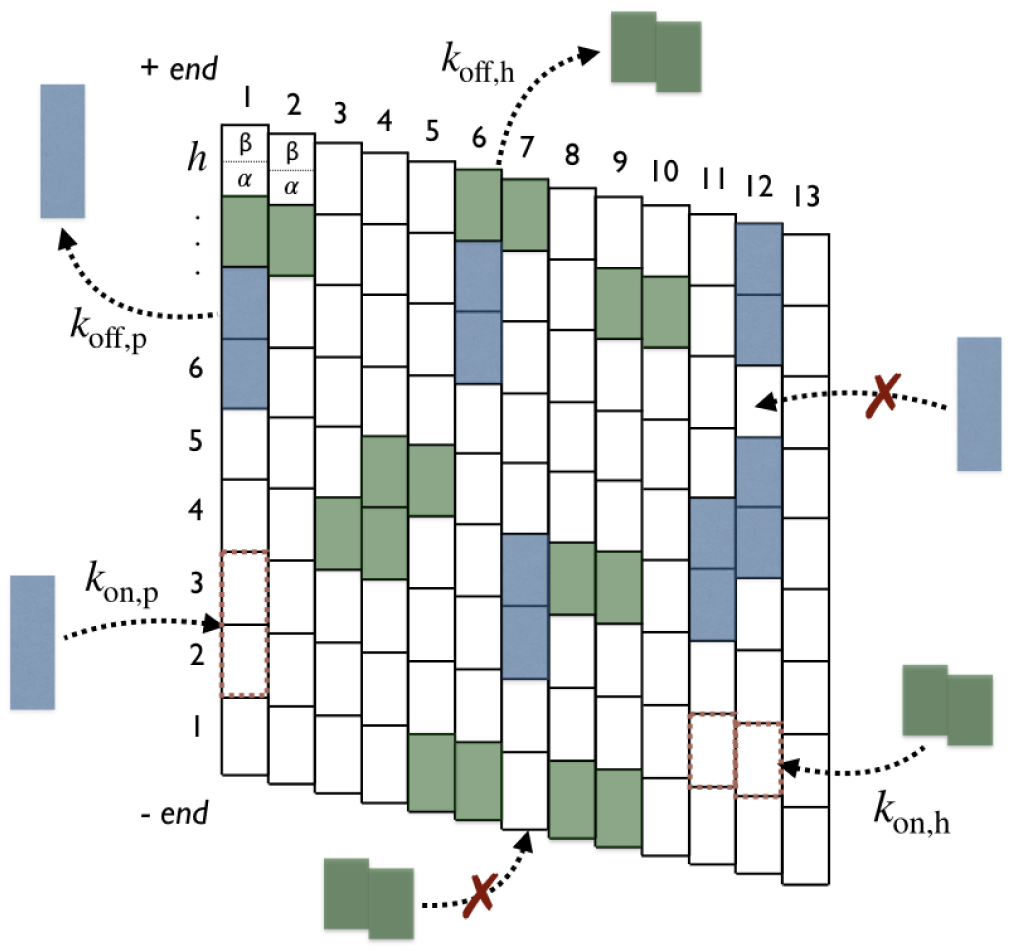
Binding rules on the microtubule lattice (*N* = 9×13 sites). Attachment and detachment of Tau’s in “*p*” (blue) and “*h*”(green) modes are represented by incoming and outgoing arrows, respectively, with respective rates: *k*_on,p_, *k*_off,p_, *k*_on,h_ and *k*_off,h_. Forbidden attachments are indicated by red crosses.

**FIG. 4:**
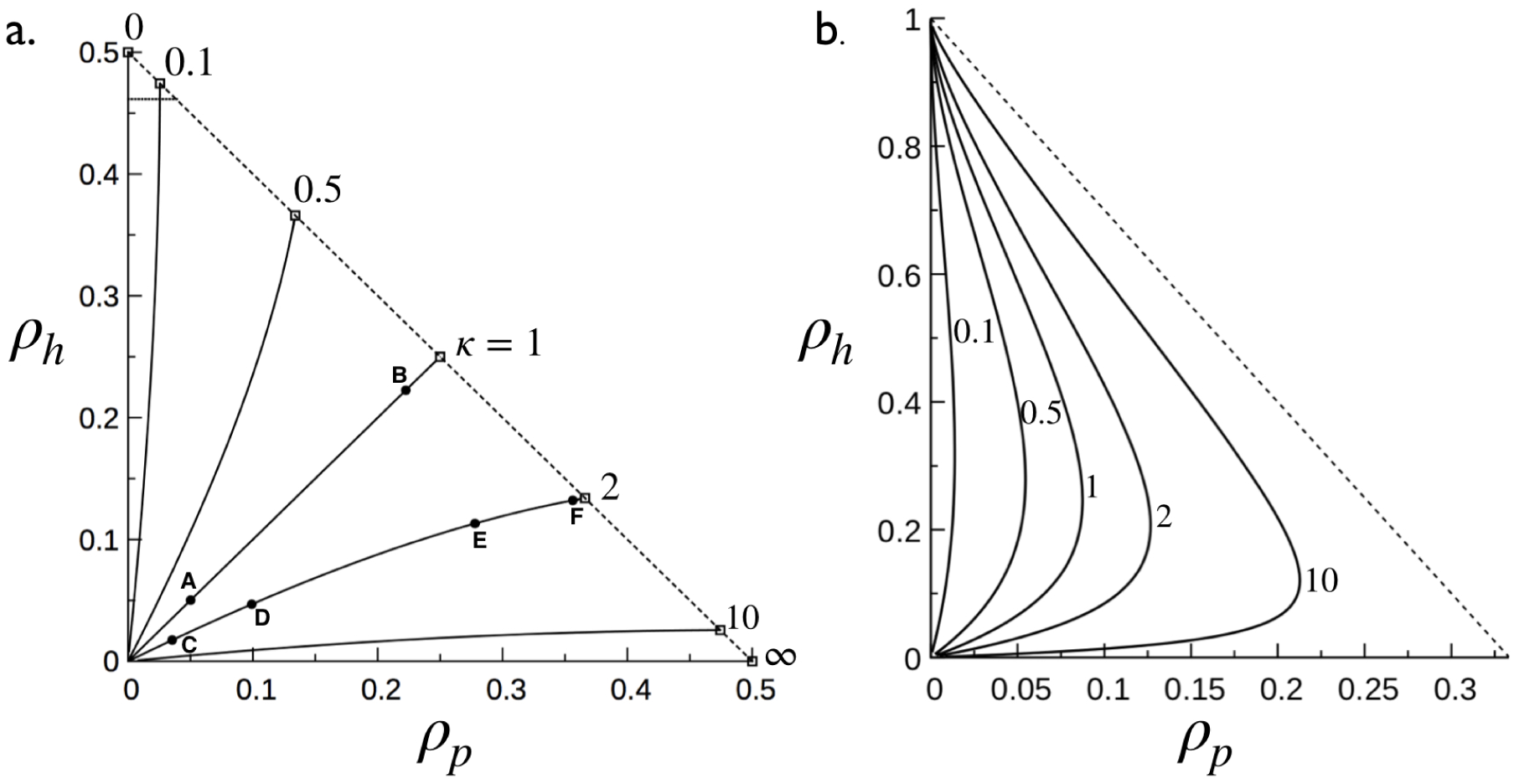
Coverage phase space of the microtubule decoration with Taus: *ρ*_*h*_ as a function of *ρ*_*p*_ for various values of *κ* (quoted numbers). Solid lines are obtained from Eq.(10) (panel **a**) and Eq.(9) (panel **b**) and dashed lines represent the saturation line given by Eq.(8) (in panel **b**, the saturation line reduces to the point (*ρ*_*p*_ = 0, *ρ*_*h*_ = 1)). Intersections between solid lines and the dashed line give the coordinates (*ρ*_*p,s*_, *ρ*_*h,s*_) at saturation. (**a**) Case of *σ*_*p*_ = *σ*_*h*_ = 1 (see Figs. 2 and 3 for illustration). Points *A* and *B* on the line *κ* = 1 correspond to *x* = 0.15 and *x* = 10, respectively, with *k*eq = 3. On the line *κ* = 2, points *C* and *D* correspond to *x* = 0.15 with *k*eq = 0.66 and *k*eq = 66, respectively, and *E* and *F* to *x* = 10 with *k*_eq_ = 0.66 and *k*_eq_ = 66, respectively. (**b**) Case of *σ*_*p*_ = 2 and *σ*_*h*_ = 0.

### Decoration kinetics

Consider a MT lattice consisting of *N* = *h* × *p* binding sites as depicted in Fig. 2. And, let *ρ*_*p*_ and *ρ*_*h*_ denotes the densities or coverages (= # bound tau / # binding sites = concentration of bound Tau / concentration of binding sites) of tau bound in mode “*p*” and “*h*”, respectively, at any time *t*. The kinetics of binding reaction is described as the “car-parking problem” (CPP) where tau-molecules reversibly bind to and detach from the MT lattice with an effective on-rate 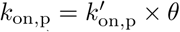 and an off-rate *k*_off,p_ for the binding “*p*” mode and with an effective on-rate 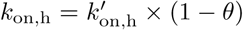 and off-rate *k*_off,h_ for the binding “*h*” mode., where *θ* is the probability of bind in “*p*” mode, 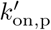 and 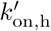 are the intrinsic on-rates for the “*p*” and “*h*” modes, respectively (see the list of parameters in Table I). In the mean field approximation, the time evolution of *ρ*_*p*_ and *ρ*_*h*_ can be described by the system of coupled non-linear differential equations:

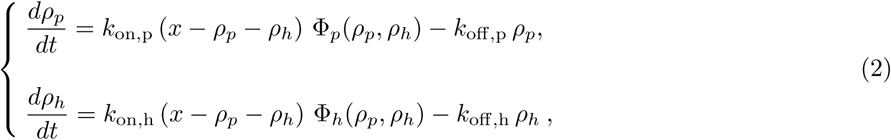

where *x* is the Tau:tubulin-dimer ratio (i.e., the ratio of the total number (concentration) of Tau-proteins in the solution over the total number (concentration) of lattice binding sites, *N* = *h* × *p*). The Φ_*p*_(*ρ*_*p*_, *ρ*_*h*_) and Φ_*h*_(*ρ*_*p*_, *ρ*_*h*_) represent the probabilities of inserting an additional tau in “*p*” and “*h*” mode, respectively, on the MT lattice already covered with a distribution of Tau at densities *ρ*_*p*_ and *ρ*_*h*_. The insertion probabilities are such that Φ_p_(*ρ*_*p*_ = 0, *ρ*_*h*_ = 0) = Φ_h_(*ρ*_*p*_ = 0, *ρ*_*h*_ = 0) = 1 for an empty MT lattice while Φ_p_(*ρ*_*p*_ = *ρ*_*p,s*_, *ρ*_*h*_ = *ρ*_*h,s*_) = Φ_h_(*ρ*_*p*_ = *ρ*_*p,s*_, *ρ*_*h*_ = *ρ*_*h,s*_) = 0 at saturation densities *ρ*_*p,s*_ and *ρ*_*h,s*_. The first terms in the right hand side of Eq.(2) represent the increasing of *ρ*_*p*_ and *ρ*_*h*_ by incorporating onto the MT lattice new tau-proteins in the respective binding mode while the second stand for the decreasing of densities as a result of the detachment of tau-proteins.

**TABLE 1:** The dynamical process of microtubule decoration with Taus is controlled by 7 key parameters. At the equilibrium, the decoration involves only 5 key parameters: *σ*_*p*_, *σ*_*h*_, *x* and the two equilibrium constants *k*_eq,p_ = *k*_on,p_*/k*_off,p_ and *k*_eq,h_ = *k*_on,h_*/k*_onh_. The main observables are the total microtubule coverage *ρ* = *ρ*_*p*_ + *ρ*_*p*_ and the nearest neighbor distribution *P*_*k*_(*r*). From these two additional observables can be derived such as the mean distance ⟨*r*_*k*_(*r*) ⟩ separating two nearest neighbor bound Taus and the order parameter *S*.

|  | Symbol | Definition |
| --- | --- | --- |
| Parameters | $\sigma_i$ | Size of Tau bound in the "$i = p, h$" mode |
| | $k_{\text{on},i}$ | Effective Tau on-rate in the "$i = p, h$" mode |
| | $k_{\text{off},i}$ | Tau off-rate in the "$i = p, h$" mode |
| | $x$ | Tau:tubulin-dimer ratio |
| Observables | $\rho_i$ | Microtubule coverage with Taos in "$i = p, h$" mode |
| | $P_k(r)$ | Nearest neighbour distribution along the direction $k = \parallel, \perp$ |
| | $\langle r_k(r) \rangle$ | Mean distance between nearest neighbors along the direction $k = \parallel, \perp$ |
| | $S$ | Order parameter |

### Observables

Now, let *ρ*_*p,eq*_ and *ρ*_*h,eq*_ denotes the equilibrium densities. For notational simplicity, we will drop in what follows the index “*eq*” on densities. Thus, the equilibrium densities *ρ*_*p*_ and *ρ*_*h*_ are obtained by setting *dρ*_*p*_*/dt* = *dρ*_*h*_*/dt* = 0 in Eq.(2). and solving the system of equations:

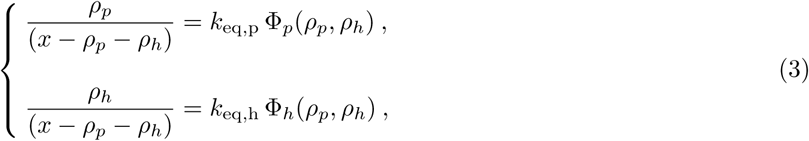

where *k*_eq,p_ = *k*_on,p_*/k*_off,p_ and *k*_eq,h_ = *k*_on,h_*/k*_off,h_ are the equilibrium constants related to the longitudinal “*p*” and the lateral “*h*” binding modes, respectively.

To characterize the two-dimensional spatial structure of *ρ*_*p*_ and *ρ*_*h*_, we consider the nearest neighbor probability distributions *P*_*‖*_ (*r*) and *P*_⊥_(*r*) of bound tau-proteins along the protofilament direction (‖) and along the helix direction (⊥), respectively, (see Fig. 1) where *r* is the unitless (in binding site unit) center-to-center distance separating two nearest-neighbors bound Tau’s, as illustrated in Fig. 5b for the protofilament or longitudinal direction (‖). The probabilities *P*_*k*=*{‖*,⊥*}*_(*r*) are given as,

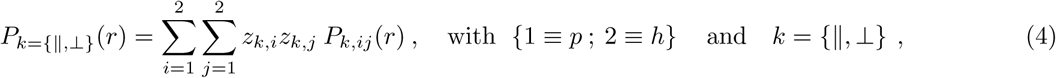

where *P*_*k,ij*_(*r*) are the partial nearest neighbour distribution (i.e., between two bound Tau’s in “*i* = *h, p*” and “*j* = *h, p*” modes) such that, 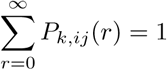, and *z*_*k,i*_, the fractions of bound Tau’s in “*i*” mode counted in the direction *k*, are given by,

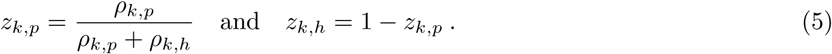

The *ρ*_*k,i*_ are the directional density (at equilibrium) of Tau bound in “*i*” mode counted in the direction *k* (see Fig. 8 for illustration and Section VI for derivation).

**FIG. 5:**
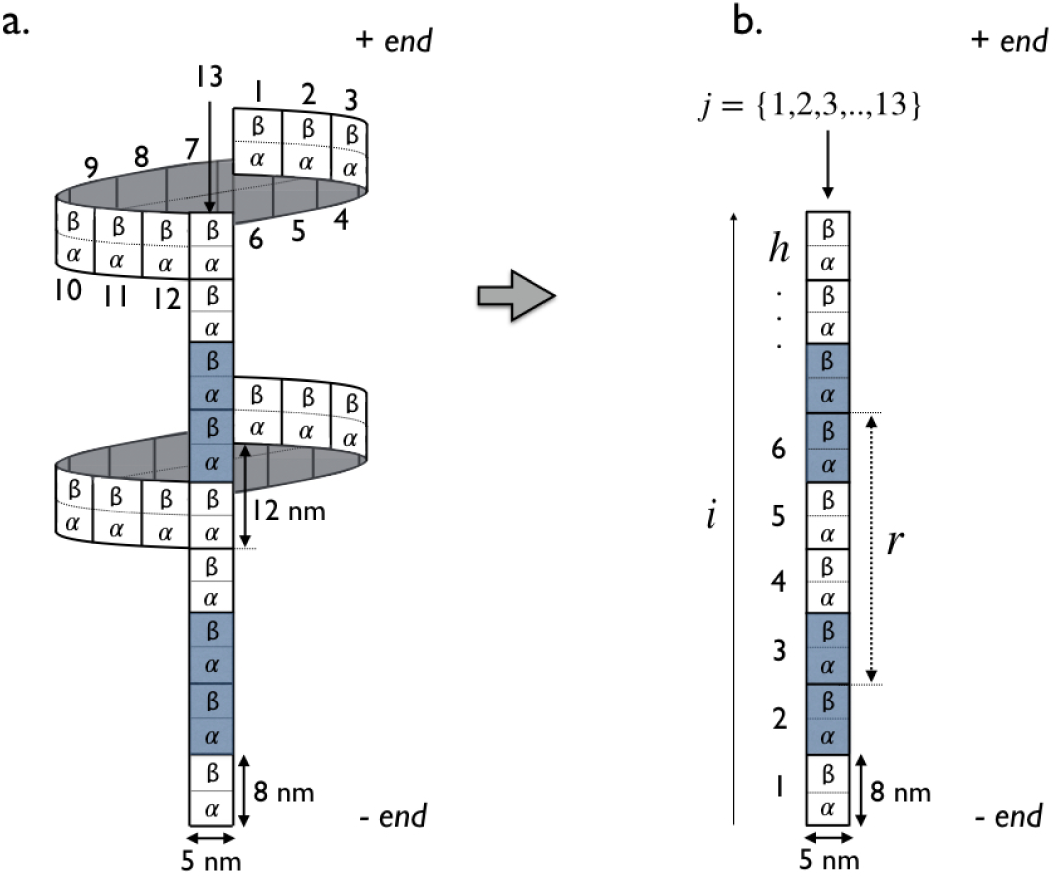
Depiction the limit case *κ*→+∞ corresponding to the single protofilament binding mode with *σp* = 1. (a) Three-dimensional representation of the 13−protofilament microtubule with two bound Taus (in blue). (**b**) In the *κ*→+∞ limit, the microtubule lattice is decoupled to consider a single protofilament consisting of *h* lattice sites (i.e., tubulin dimers). The spatial extension of a lattice site corresponds to the *αβ*-tubulin dimer length of 8 *nm*. The unitless center-to-center distance between two nearest neighbors bound-tau is denoted by *r* and equals to *r* = 4 in this example (equivalent to 32 *nm*).

To have a global picture characterizing the overall trend of Tau spatial arrangements on the MT lattice, we define the order parameter *S* (such as, −1 ≤ *S* ≤ +1) as,

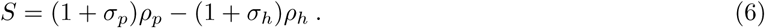

*S* = +1 describes the case where all bound Tau’s are aligned along the protofilaments whereas *S* = ‒ 1 the one where the bound Tau’s are all aligned along the MT helices and *S* = 0 corresponds to the 50 ‒ 50 situation.

In summary, the decoration of a MT with Tau will be characterized by the following observables (at equilibrium):

- *ρ* = *ρ*_*p*_ + *ρ*_*h*_: the (longitudinal, lateral and) total MT coverage by Tau-proteins,
- *P*_*k*=*{*‖,⊥*}*_(*r*): the averaged distributions (longitudinal *k* =‖ and lateral *k* =⊥) of spacing between bound Tau-proteins at the MT surface,
- *S*: the order parameter.

In what follows, we will the determine expressions for these observables and analyze their changes as functions of key parameters (in Table I) controlling the decoration kinetics.

## IV. RESULTS

### A. General results

The key functions in the dynamical microtubule decoration process are the probabilities Φ_p_ and Φ_h_ of inserting an additional Tau-molecule in “*p*” and “*h*” modes, respectively, on the microtubule already covered with a distribution of Tau’s at densities (*ρ*_*p*_, *ρ*_*h*_). For the system under consideration (binding rules in Sec. III and illustration in Fig. 3) the Φ_p_ and Φ_h_ for noncooperative binding of Tau-molecules are given by (see Sec. VI for the derivation):

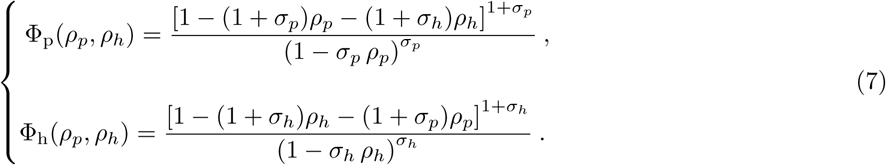

By construction, Φ_i_(*ρ*_*p*_, *ρ*_*h*_) (*i* = *p, h*) satisfy the criteria: Φ_i_(*ρ*_*p*_ = 0, *ρ*_*h*_ = 0) = 1 for an empty MT and Φ_i_(*ρ*_*p,s*_, *ρ*_*h,s*_) = 0 at the saturation densities (*ρ*_*p,s*_, *ρ*_*h,s*_) when all binding sites on the MT are occupied by Tau-molecules. (*ρ*_*p,s*_, *ρ*_*h,s*_) can be obtained by equating the numerators of Φ_p_ and Φ_h_ above to zero to give an ensemble of (*ρ*_*p,s*_, *ρ*_*h,s*_) belonging to the line:

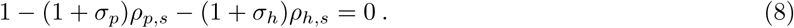

As we are dealing with more than one kind of ligands (Tau binding modes), an additional condition to Eq.(8) is required to determine specific values of (*ρ*_*p,s*_ and *ρ*_*h,s*_).

### 1. Microtubule coverage

#### Phase space

In general, analytical solutions of Eq.(3) providing expressions for *ρ*_*p*_ and *ρ*_*h*_ are not possible because of the non-linearities in the probabilities Φ_p_ and Φ_h_, However, some insights can already be gained by analyzing the phase space (*ρ*_*p*_, *ρ*_*h*_) without having solved Eq.(3) (i.e., determining *ρ*_*p*_ and *ρ*_*h*_).

In the (*ρ*_*p*_, *ρ*_*h*_) space, the physical space for possible values of *ρ*_*p*_ and *ρ*_*h*_ [(0, 0) ≤ (*ρ*_*p*_, *ρ*_*h*_) ≤ (*ρ*_*p,s*_, *ρ*_*h,s*_)] is a rectangle triangle delimited by the horizontal *ρ*_*h*_ = 0 and vertical *ρ*_*p*_ = 0 axes and the saturation line, 1 ‒ (1 + *σ*_*p*_)*ρ*_*p*_ ‒ (1 + *σ*_*h*_)*ρ*_*h*_ = 0, that originates from positivity condition of insertion probabilities, Φ_p_ ≥ 0 and Φ_h_ ≥ 0. Using expressions in Eq.(7) back into Eq.(3) leads to the ratio for the densities, equation

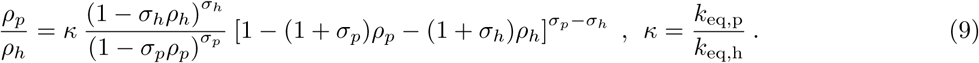

Now, solving Eq.(9) allows us to portray *ρ*_*h*_ as a function of *ρ*_*p*_ within the triangular section of the phase space as shown in Fig. 4, wherein *ρ*_*p*_ and *ρ*_*h*_ are solutions of Eq.(3). Eq.(9) shows that *κ* is the key organizing parameter of portraits *ρ*_*h*_ *vs ρ*_*p*_ (parameterized by the *σ*’s) that all terminate on the saturation line, yielding the values *ρ*_*p,s*_ and *ρ*_*h,s*_ (see, Fig. 4). Therefore, combining Eqs.(8) and (9) allows determining *ρ*_*p,s*_ and *ρ*_*h,s*_. As analytical solutions of Eq.(9) are only possible for *σ*_*i*_ = 0, 1, we consider without loss of generality the cases displayed in Fig. 4 for which the solution of Eq.(9) is,

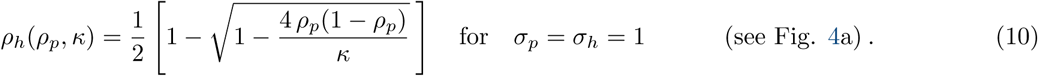

When *σ*_*p*_ = *σ*_*h*_, Eq.(9) shows that *ρ*_*h*_ = *ρ*_*p*_ is the trivial solution for *κ* = 1 and that portraits of *ρ*_*h*_ *vs ρ*_*p*_ for *κ* ≠ 1 are symmetric about the *κ* = 1 trajectory, i.e., *ρ*_*h*_(*ρ*_*p*_, *κ*) = *ρ*_*p*_(*ρ*_*h*_, 1*/κ*), as illustrated in Fig. 4a for *σ*_*p*_ = *σ*_*h*_ = 1. In addition to explicitly depending on *κ*, positions along the trajectory *ρ*_*h*_ *vs ρ*_*p*_ depend on the kinetic parameters: the tau:tubulin-dimer ratio, *x*, and the effective equilibrium constant, *k*_eq_ = *k*_eq,p_ + *k*_eq,h_. For example, points *A* and *B* on the trajectory *ρ*_*h*_(*ρ*_*p*_, *κ* = 1) = *ρ*_*p*_ correspond to identical, *k*_eq,p_ = *k*_eq,h_ = 1.5 (i.e., *k*_eq_ = 3) but to different ratios, *x* = 0.15 and *x* = 10, respectively. This indicates that for a given set of (*k*_eq,p_, *k*_eq,h_), the trajectory *ρ*_*h*_(*ρ*_*p*_, *κ*) approaches the saturation line with increasing *x*. Using the saturation line [*], 1 ‒ 2*ρ*_*p,s*_ − 2*ρ*_*h,s*_ = 0 (for *σ*_*p*_ = *σ*_*h*_ = 1), back into Eq.(9) and solving for *ρ*_*p,s*_ and *ρ*_*h,s*_ leads to:

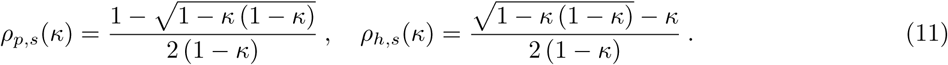

Note that the partial saturation densities *ρ*_*p,s*_ and *ρ*_*h,s*_ are functions of *κ* but the total density, *ρ*_*s*_ = *ρ*_*p,s*_ + *ρ*_*h,s*_ = 1/2, is not as it should be.

When *σ*_*p*_ ≠ *σ*_*h*_, the general trend of portraits *ρ*_*h*_ *vs ρ*_*p*_ is quite different from that of *σ*_*p*_ = *σ*_*h*_ in the sense that *ρ*_*h*_ as a function of *ρ*_*p*_ is now bi-valued with an extremum and all portrait lines converge to the saturation coordinates (*ρ*_*p,s*_ = 0, *ρ*_*h,s*_ = 1/(1 + *σ*_*h*_)) for *σ*_*p*_ *> σ*_*h*_ or (*ρ*_*p,s*_ = 1/(1 + *σ*_*p*_), *ρ*_*h,s*_ = 0) for *σ*_*p*_ *< σ*_*h*_, i.e., the system converges to the highest stoichiometry at the saturation. Fig. 4b illustrates the case of *σ*_*p*_ = 2 and *σ*_*h*_ = 0 that leads to solving a cubic equation to determine *ρ*_*p*_ and *ρ*_*h*_.

#### Diluted regime

The diluted regime, corresponding to a low MT coverage *ρ*_*p*_≪ 1 and *ρ*_*h*_≪ 1, is both especially relevant to the context of axons [28] and interesting as it allows deriving analytical solutions of Eq.(3). In this case, the insertion probabilities in Eq.(7) can be linearized to the leading order as,

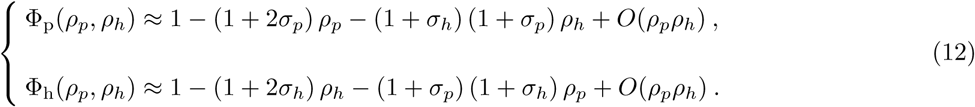

In addition, the trajectories *ρ*_*h*_ *vs ρ*_*p*_, given in Eq.(9) and illustrated in Fig. 4, can also be linearized as, *ρ*_*h*_ ≃*ρ*_*p*_*/κ* + *O*(*ρ*_*p*_^2^). Now, using Eq.(12) back into Eq.(3) and with linearized phase portraits leads to a system of equations that can be solved analytically to give,

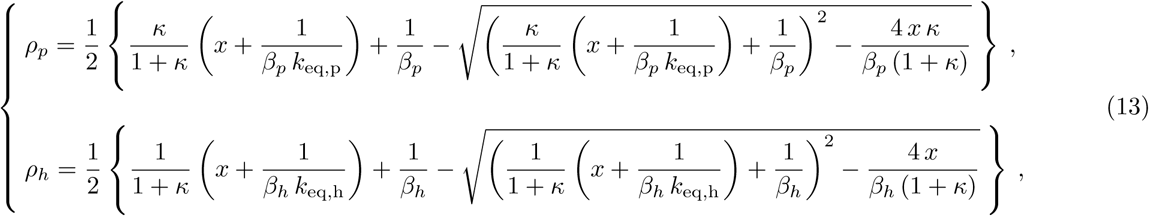

where the coefficients *β*_*p*_ and *β*_*h*_ are given by

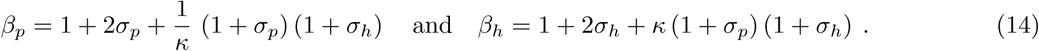

For low tau:tubulin-dimer ratio, *x* ≪ 1, the densities become linear functions of *x* as,

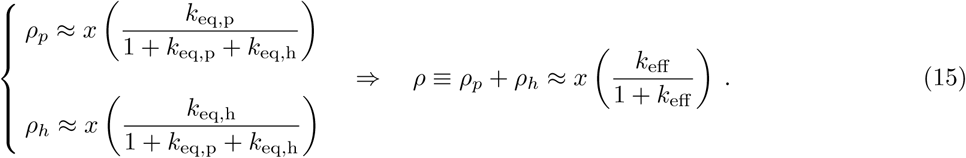

Interestingly, in the limit *x* ≪1, the coverage is independent of the binding reaction stoichiometries and, therefore, fitting the experimental data (as, for example, in cosedimentation assays) in this regime would allow determination of the effective equilibrium constant *k*_eff_ = *k*_eq,h_ + *k*_eq,p_. Out of the *x* ≪ 1 regime, the *ρ*’s given in Eq.(13) are non-linear functions of *x* and depend on stoichiometries.

### 2. Distribution of Tau-proteins on MT surface

From an experimental point of view, it turns out quite challenging to resolve the distribution of Tau-proteins along the MT helices, i.e., the lateral distribution *P*_⊥_(*r*). Therefore, we will only discuss the longitudinal properties *P*_‖_ (*r*) that can be investigated from experimental data [38] and leave *P*_⊥_(*r*) in the Method Section VI.

We recall that *P*_‖_ (*r*) is the probability distributions, along the protofilament direction (‖), of nearest neighbor Tau-proteins bound on the MT surface. *P*_‖_(*r*) is obtained from Eq.(4) where *z*_‖,*p*_ = *ρ*_*p*_/ [*ρ*_*p*_ + (1 + *σ*_*h*_)*ρ*_*h*_], *z*_‖,*h*_ = 1 *z*_‖,*p*_, and the partial distributions *P*_‖,*ij*_(*r*) between two Tau’s bound in “*i*” and “*j*” modes given by (see Section VI for derivations)

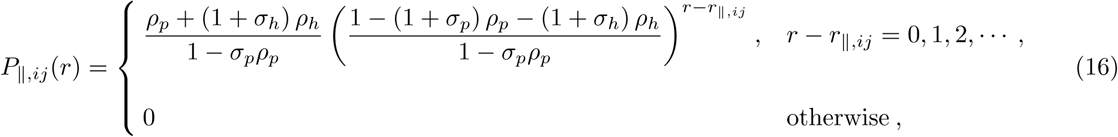

where *r*_‖,*ij*_ is the element of the matrix *r*_‖_ of the minimum physical center-to-center distances between two neighbors Tau’s (bond in “*i*” and “*j*” modes) along the protofilament,

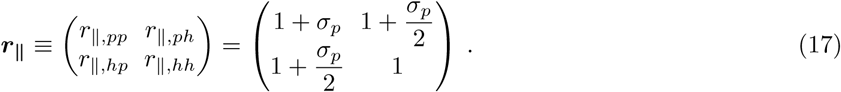

For instance, the minimum center-to-center between two Tau’s bound in “*p*” and “*h*” modes is *r*_‖,*ph*_ = *r*_‖,*hp*_ = [(1 + *σ*_*p*_) + 1] /2 since a Tau bound in “*p*” mode covers 1 + *σ*_*p*_ binding sites along the protofilament and the one bound in “*h*” mode covers 1 binding site along the protofilament. The resulting distribution *P*_‖_ (*r*) of Tau-proteins on MT surface is multi-exponential with the mean distance separating two nearest neighbor bound Tau along protofilaments given by,

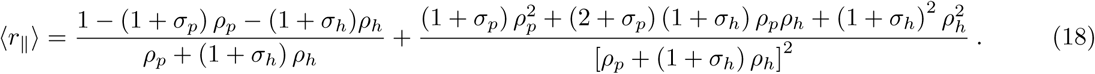

Note that, 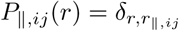 and ⟨*r*_‖_ ⟩ = 1/ (1 ‒ *σ*_*p*_*ρ*_*p,s*_), at the saturation limit, 1‒ (1 + *σ*_*p*_) *ρ*_*p,s*_‒ (1+*σ*_*h*_)*ρ*_*h,s*_ = 0.

We now consider specific cases to gain more insights into the MT decoration.

### B. Single binding modes

In the case of a single binding mode “*i*” the MT decoration is controlled by 3 key parameters (see Table I): the Tau binding size *σ*_*i*_ (or the binding stoichiometry *ν*_*i*_ = 1/(1 + *σ*_*i*_)), the equilibrium constant *k*_eq,i_ and the Tau:tubulin-dimer ratio *x*.

#### 1. κ → +∞: protofilament binding mode

The *κ* → + ∞ limit corresponds to *k*_eq,h_ = 0 meaning that the “*p*” mode is the only possible mode of binding for Tau’s (see Fig. 5). The MT decoration is characterized as follows:

- *MT coverage:* In this limit, *ρ*_*h*_ = 0 (corresponding to the x-axis in the phase space in Fig. 4a) and the total coverage, *ρ* = *ρ*_*p*_, lies between 0 and the saturation *ρ*_*s*_ = 1/(1 + *σ*_*p*_) ≡*ν*_*pp*_, depending of *k*_eq,p_ and *x*. Figure 6a shows numerical solutions (solid lines) of Eq.(3) along with simulations results (data points) of *ρ* as a function of *x* for *σ*_*p*_ = 1 and various *k*_eq,p_. At low *x*, the coverage *ρ* linearly increases with *x* as predicted in Eq.(15), then deviates from linearity and logarithmically reaches the saturation *ρ*_*s*_ ≡*ν*_*pp*_ = 0.5 at high *x*. In the dilute regime, the coverage *ρ* is obtained by taking the *κ* → + ∞ limit in Eq.(13) to give:

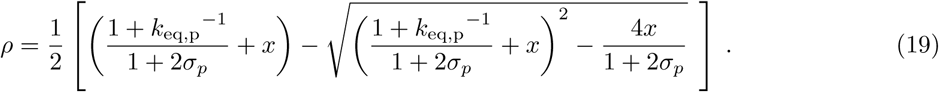
- *Distribution of Tau spacing:* The nearest neighbor distribution, *P*_‖_(*r*), and its first moment, ⟨*r*_‖_⟩, are obtained by taking the *κ* → +∞ limit in Eqs.(4), (16) and (18) to obtain:

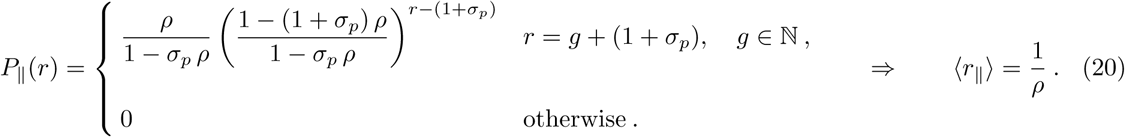 The most likely spacing between Taus along the protofilament axis is *r*_‖,*max*_ = (1 + *σ*_*p*_), corresponding to *σ*_*p*_ = 1 to *r*_‖,*max*_ = 2 × 8 *nm* = 16 *nm* (in real units), for the case *σ*_*p*_ = 1 illustrated in Fig. 5, while the mean Tau spacing, ⟨*r*_‖_⟩ = 1*/ρ*, range between 16 *nm* (for *ρ* ≡ *ρ*_*s*_ = 1/2 at the saturation) and (8*/ρ*) *nm*. For example, for the minimal coverage, *ρ* = 2*/h*, of two Taus per protofilament of length *h*, we have (8*/ρ*) *nm* = 4*h nm* and for a typical microtubule of length 5 *µm*, corresponding to *h* = 5 *µm/*8 *nm* = 625, we found (8*/ρ*) *nm* = 2.5 *µm*.
- *Spatial arrangement of Taus:* To illustrate the spatial arrangement of Taus on the MT surface, we consider two contrasted configurations at low (*x* = 0.15, *ρ* ≈ 0.1) and high (*x* = 10, *ρ* ≈ 0.45) densities (Tau:tubulin-dimer ratio and coverage) at the same equilibrium constant *k*_eq,p_ = 3. These two configurations are indicated by filled circles in Fig. 6a and snapshots of Tau arrangement on MT surface with associated Tau spacing distribution *P*_‖_(*r*) are displayed in Figs. 6b-e (snapshots and histogram in *P*_‖_(*r*) are from simulations and lines are from Eq.(20)). At low density, corresponding to an order parameter *S* ≈ 0.2, there is no apparent spatial organization of Taus (Fig. 6b) and their spacing distribution is an exponential decay (Fig. 6c) with a mode probability ≈10% of finding a longitudinal spacing of 16 *nm*. In contrast, at density with *S*≈ 0.9, there is a spatial order in the protofilament direction (Fig. 6e) and the nearest neighbor spacing distribution in Fig. 6d shows a sharp exponential decay with a maximum probability ≈ 85% at spacing spacing of 16 *nm*. At the saturation limit, *ρ* = 1/2 and *S* = 1; all the Tau-proteins are perfectly aligned along the protofilament with a nearest neighbor distribution given by a Dirac delta function centered on *r* = (1 + *σ*_*p*_) = 2 (16 *nm* in real units). By analogy with crystalline liquids, the configurations of Fig. 6b can be defined as a nematic-type phase, whereas those of Fig. 6e are defined as a smectic-type phase.

**FIG. 6:**
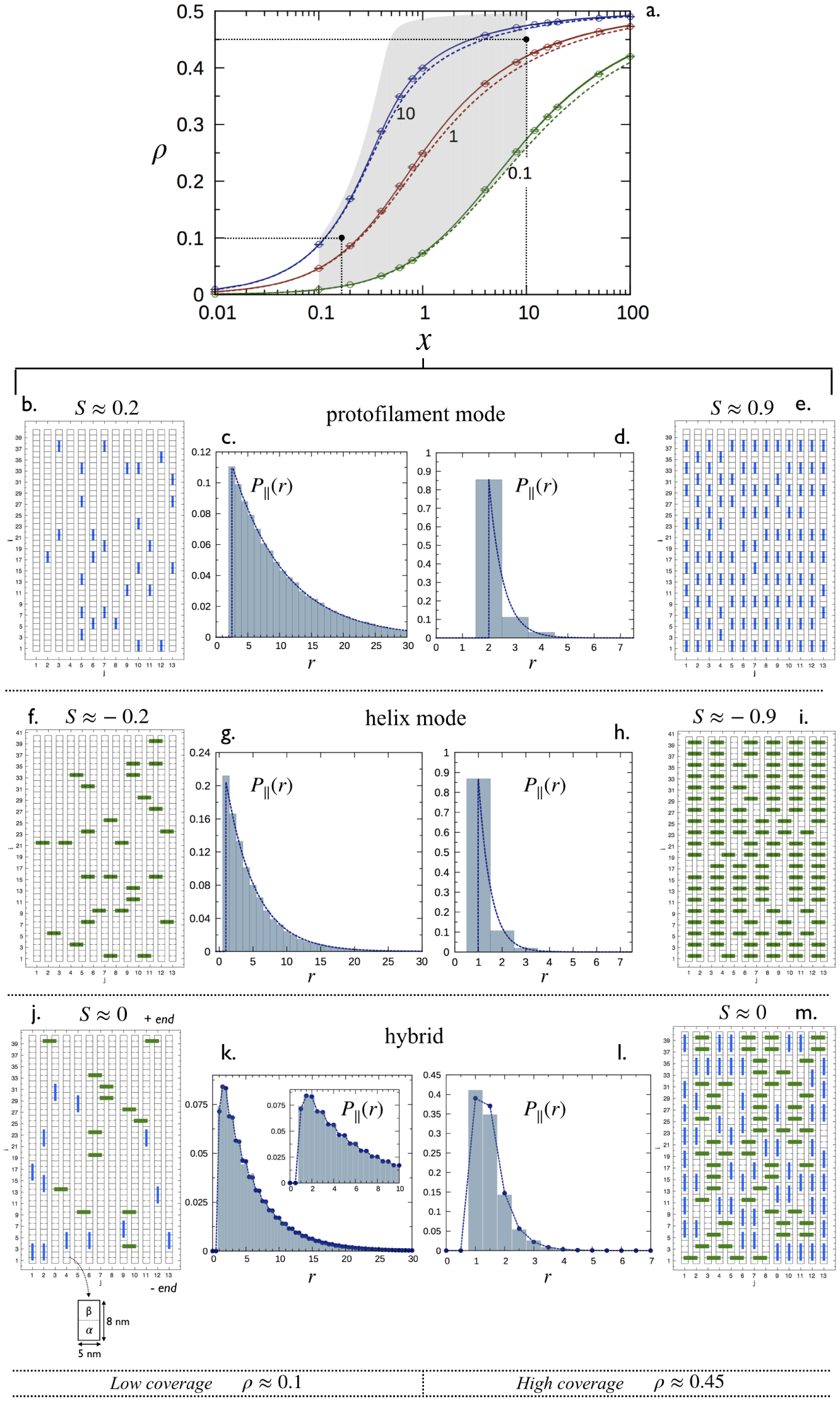
Decoration of a microtubule by a population of tau proteins. (**a**) Saturation curves: total coverage *ρ* at equilibrium state as a function of *x* for *σ*_*p*_ = *σ*_*h*_ = 1. Point data (circles) represent results from Monte Carlo simulations in the limit case *κ* → +∞ (mode “*p*”) with *k*eq,p = 0.1, 1 and 10. Solid lines correspond to numerical solutions for *ρ* of Eq. (3) in the limit case *κ* → +∞ while dashed lines correspond to numerical resolution of (3) in the limit case *κ* = 1 (see Sec. VI). The grayed zone gives the domain of coverage consistent with the axonal conditions. The zone has been obtained using the two ranges 0.1 ≤ *x* ≤ 10 and 0.1 ≤ *k*_eff_ ≤ 10^3^ (see VI for the estimation of these ranges from the literature). The two points correspond to *A* and *B* in Fig. 6. (**b**-**m**) Snapshots and their corresponding distributions *P*_‖_ (*r*) for one binding mode in (**b**-**e**), (**f** -**i**) and two binding modes with *κ* = 1 in (**j**-**m**). In each case, the decoration is characterized for low *ρ* ≈ 0.1 (*x* = 0.15), and high coverages, *ρ* ≈ 0.45 (*x* = 10). Histograms for the nearest neighbour distributions in (**c**,**d**,**g**,**h**,**k**,**l**) have been calculated from Monte Carlo simulations (see section method VI). Dashed lines in (**c**,**d**,**g**,**h**,**k**,**l**) correspond to theoretical distributions obtained using Eq. (20) for the mode “*p*” in (**c**,**d**), Eq. (21) for the mode “*h*” in (**g**,**h**) and Eqs. (16), (4) for the two binding modes in (**k**,**l**).

#### 2. κ → 0: helix binding mode

The *κ* → 0 limit corresponds to *k*_eq,p_ = 0 implying that Taus can bind on the MT surface only in the “*h*” mode. This case is very similar to the *κ* → +∞ limit discussed above. Indeed:

- *MT coverage:* In this limit, *ρ*_*p*_ = 0 (corresponding to the y-axis in the phase space in Fig. 4a) and the total coverage, *ρ* = *ρ*_*h*_, lies between 0 and the saturation *ρ*_*s*_ = 1/(1 + *σ*_*p*_*h*) ≡*ν*_*hh*_, depending on *k*_eq,h_ and *x*. The variations of *ρ* as a function of the key parameters *k*_eq,h_ and *x* are exactly the same to that of *κ* → + ∞ limit. Therefore, the two limits *κ* → + ∞ and *κ* → 0 are indistinguishable in terms of microtubule coverage as we are dealing with a single binding mode. In the dilute regime, the *κ* → 0 limit of Eq.(13) leads the same expression to the one obtained in the *κ* → + ∞ limit by replacing the subscript “*p*” by “*h*” in Eq. (19).
- *Distribution of Tau spacing:* Taking the *κ* → 0 limit Eqs.(4), (16) and (18) gives:

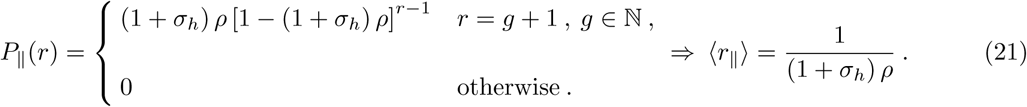 Note that this expression is different from the one in Eq.(20). As a consequence, the most likely spacing between Taus along the protofilament axis is *r*_‖,*max*_ = 1 (i.e., *r*_‖,*max*_ = 8 *nm* (in real units)) while for *σ*_*h*_ = 1 the mean Tau spacing, ⟨*r*_‖_ ⟩= 1/(2*ρ*), range between 8 *nm* (for *ρ* ≡ *ρ*_*s*_ = 1/2 at the saturation) and (4*/ρ*) *nm*. Likewise for the minimal coverage, *ρ* = 2*/h*, of two Taus per protofilament of length *h*, we have (4*/ρ*) *nm* = 2*h nm* and for a typical microtubule of length 5 *µm*, corresponding to *h* = 5 *µm/*8 *nm* = 625, we found (4*/ρ*) *nm* = 1.25 *µm*.
- *Spatial arrangement of Taus:* As in the *κ* → +∞ limit, Figs. 6f-i display snapshots of Tau arrangement on MT surface with associated Tau spacing distribution *P*_‖_(*r*) for the same low and high density configurations in the *κ* → 0 limit (snapshots and histogram in *P*_‖_ (*r*) are from simulations and lines are from Eq.(21)). Apart from the order parameters that are now negative, *S* ≈ ‒0.2 and *S* ≈ ‒0.9 for low and high density, respectively, all the observations made in the *κ* → +∞ limit also prevail for the *κ* → 0 limit.

### C. Two binding modes

When both two Tau binding modes can occur, the MT decoration involves 5 key parameters (see Table I): two Tau binding sizes *σ*_*p*_ and *σ*_*h*_, two equilibrium constants *k*_eq,p_ and *k*_eq,h_, and the Tau:tubulin-dimer ratio *x*. It thus follows that 4 contrasted situations can be distinguished: *κ* = 1 with *σ*_*p*_ = *σ*_*h*_ and *κ* ≠ 1 with *σ*_*p*_ = *σ*_*h*_ and *σ*_*p*_≠ *σ*_*h*_.

#### 1. κ = 1: identical equilibrium constants

- *MT coverage:* The phase spaces in Fig. 4 show two different trajectories of *κ* = 1 (in panels *a* and *b*). As can be seen, and already emphasized in the text below Eqs.(9) and (10), the portraits are linear, *ρ*_*h*_ = *ρ*_*p*_ for *σ*_*p*_ = *σ*_*h*_, and non-linear otherwise, and positions, (*ρ*_*p*_, *ρ*_*h*_), along that trajectories depends on both the effective equilibrium constant, *k*_eff_ = *k*_eq,p_ + *k*_eq,h_, and the Tau:tubulin-dimer ratio *x*. Numerical results for the total coverage *ρ* as a function of *x*, for *σ*_*p*_ = *σ*_*h*_ = 1 and three values of *k*_eff_ = 0.1, 1 and 10, are shown by dashed lines in Fig. 6a. Clearly, the behaviors of *ρ* as a function of *x* are very similar in both single (*κ* → + ∞ and *κ* → 0 limits) and two binding modes. However, the situation is quite different when *σ*_*p*_ ≠*σ*_*h*_ (see supplementary information, Fig. S1). Indeed, Fig. 4b for *κ* = 1 and *σ*_*p*_ = 2 and *σ*_*h*_ = 0 shows that *ρ*_*h*_ monotonically increases from zero to saturation around *ρ*_*h*_ = 1 while *ρ*_*p*_ (< *ρ*_*h*_) increases from zero reaches a maximum and decreases when approaching saturation conditions. This indicates that for any *k*_eff_ and *x* below the saturation, the system is bi-phasic and admits two equilibrium coverages with identical *ρ*_*p*_ and different *ρ*_*h*_. At saturation, the system becomes mono-phasic involving only the binding mode with smaller *σ*.
- *Distribution of Tau spacing:* Inspection of Figs. 6j-m (snapshots and histogram in *P*_‖_(*r*) are from simulations and lines are from Eqs.(4) and (16)) shows that allowing two binding modes for Taus effectively impact the spacing distributions. As can be seen in Figs. 6k,l, *P*_‖_(*r*) in Eq.(4) is no longer a single exponential distribution but rather a summation of exponential decays with *P*_‖,*ij*_(*r*) given in Eq.(16). The difference in the *P*_‖_(*r*) shape, in comparison with that of the single binding modes, is particularly noticeable for configurations closed to the saturation, as illustrated in Fig. 6l.
- *Spatial arrangement of Taus:* The low (*x* = 0.15, *ρ* ≈ 0.1) and high (*x* = 10, *ρ* ≈ 0.45) density configurations considered above in single binding modes correspond here to the points *A* and *B* (with *σ*_*p*_ = *σ*_*h*_ = 1 and *k*_eff_ = 3) along the linear trajectory *ρ*_*h*_ = *ρ*_*p*_ in the phase space in Fig. 4a. As *σ*_*p*_ = *σ*_*h*_, both configurations *A* and *B* are characterized by an order parameter *S* ≈ 0, i.e., there is on average the same amount of Taus bound in “*p*” and “*h*” modes as can be seen in the snapshots of Figs. 6j and m. However, as shown in Fig. 4b for *κ* = 1 and *σ*_*p*_ = 2 and *σ*_*h*_ = 0 the curve *κ* = 1 is can be found below or above the line *ρ*_*h*_ = 3*ρ*_*p*_ depending on *k*_eff_ and *x*. Therefore, there exist threshold values of *k*_eff_ and *x* at which the order parameter *S* = 0, below and above which it is *S* > 0 and *S* < 0 respectively. At the saturation, *S* < 0 for *σ*_*p*_ > *σ*_*h*_ and vice versa.

#### 2. κ ≠ 1: non-identical equilibrium constants

To illustrate the MT decoration in the situation of non-identical equilibrium constants of binding modes, we consider the of case *κ* = 2, shown in the phase space in Fig. 4a for *σ*_*p*_ = *σ*_*h*_ = 1. The 4 points along the that trajectory correspond to the following configurations: *C* = (*x* = 0.15, *ρ* ≈ 0.05, *k*_eff_ = 0.66) and *D* = (*x* = 0.15, *ρ* ≈ 0.15, *k*_eff_ = 66) for low density, and *E* = (*x* = 10, *ρ* ≈ 0.39, *k*_eff_ = 0.66) and *F* = (*x* = 10, *ρ* ≈ 0.48, *k*_eff_ = 66) for high density.

- *MT coverage:* Figs. 7a and b show the partial and total coverage as a function of *x*, for *σ*_*p*_ = *σ*_*h*_ = 1 and *k*_eff_ = 0.66 and 66. Configurations *C* and *E* are represented by the two filled circle points on *ρ vs x* in Fig. 7a, and *D* and *F* by the filled circle points on the same curve in Fig. 7b.Densities *ρ*_*p*_ and *ρ*_*h*_ linearly increase with *x* at low *x* and logarithmically reach their saturation 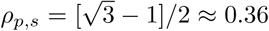 and 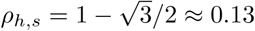 (as predicted in Eq.(11)) at high *x*. In any cases, we have *ρ*_*p*_ > *ρ*_*h*_.

**FIG. 7:**
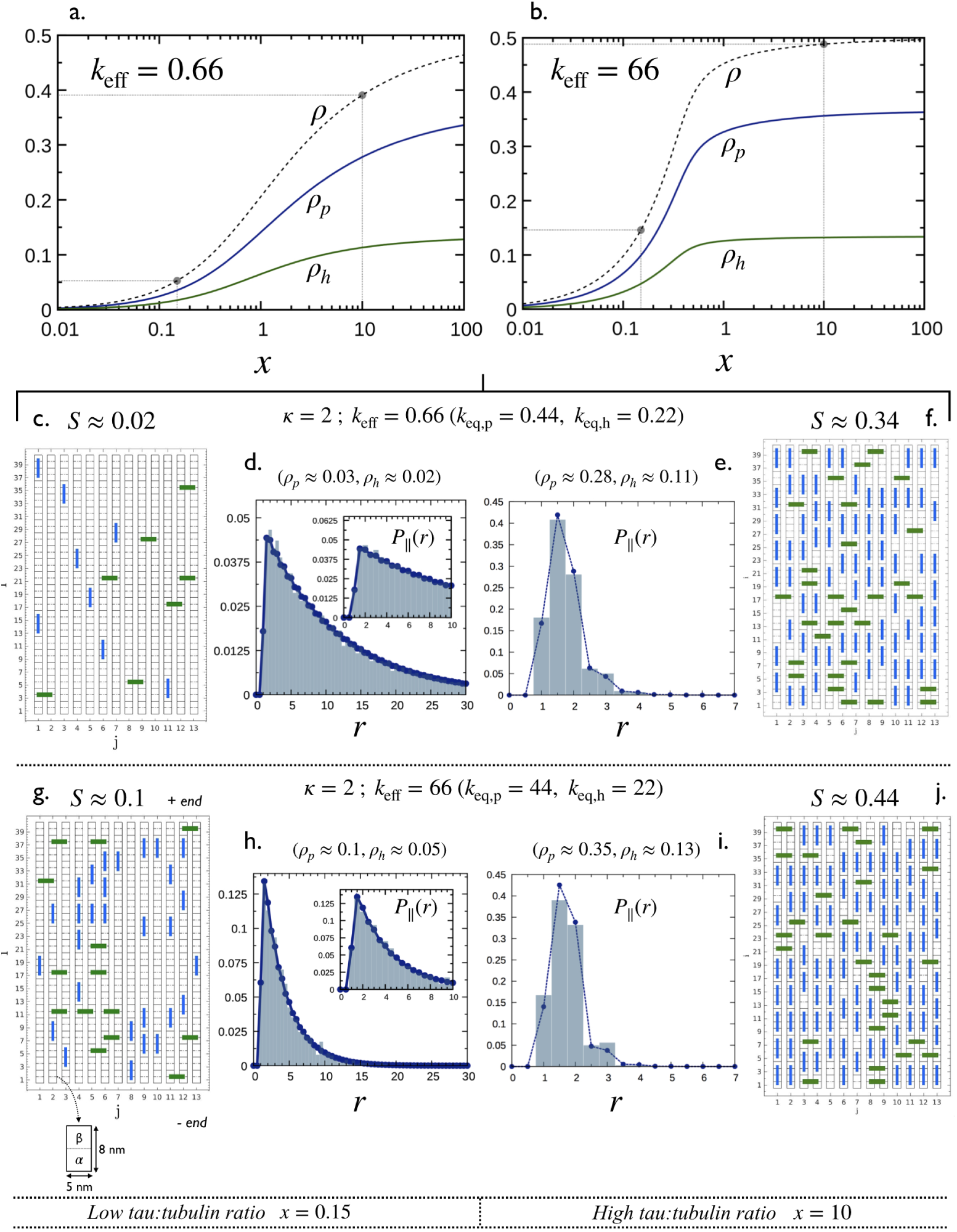
Decoration of a microtubule by tau for the case *κ* = 2 (i.e., *k*_eq,p_ = 2 *k*_eq,h_) and *σ*_*p*_ = *σ*_*h*_ = 1. (**a**,**b**) Densities *ρ*_*p*_, *ρ*_*h*_ and the total coverage *ρ* = *ρ*_*p*_ + *ρ*_*h*_ as a function of the ratio, *x* for an effective equilibrium constant *k*_eff_ = 0.66 and 66, respectively. Solid lines correspond to numerical resolution of Eq. (3). In both cases, results corresponding to *x* = 0.15 and *x* = 10 are highlighted in black point circles. This leads to 4 configurations: *C, D, E* and *F* (see Fig. 4b) for which typical snapshots and averaged longitudinal spacing distributions, *P*_‖_(*r*) are shown in (**c**-**j**). Histograms in (**d**,**e**,**h**,**i**) have been calculated from Monte Carlo simulations (see section method VI) while dashed lines with blue points have been obtained using Eqs. (16) and (4).

- *Distribution of Tau spacing:* Snapshots of MT decoration and associated nearest neighbor distributions for the 4 configurations *C, D, E* and *F* (filled circle points in Figs. 7a and b) are shown in Fig. 7c-j (snapshots and histogram in *P*_‖_(*r*) are from simulations and lines are from Eqs.(4) and (16)). The shape of the distributions in Fig. 7d and h, corresponding to the low density configurations *C* and *E*, are very similar to that in Fig. 6k for *κ* = 1 (i.e., configuration *A* in Fig. 4b) with a multi-exponentional decay, while the shape for high density configurations *D* and *F* exhibit significant differences. Indeed, the main peak of *P*_‖_(*r*) in Fig. 7e and i is centered at *r* = 1.5, corresponding to close packing between Taus bound in longitudinal “*p*” (in blue) and lateral “*h*” (in green) modes. In addition, the peak centered at *r* = 2, corresponding to close packing between Taus bound in “*p*” mode, is higher than that centered at *r* = 1 for Taus bound in “*h*” mode. This is because for *κ* = 2 there are more Taus bound in “*p*” mode than those bound in “*h*” modes, i.e, *ρ*_*p*_ > *ρ*_*h*_.
- *Spatial arrangement of Taus:* As *ρ*_*p*_ > *ρ*_*h*_ and that the curve *κ* = 2 in Fig. 4b is below the line *ρ*_*h*_ = *ρ*_*p*_, all configurations in this case are characterized by an order parameter *S* > 0 as shown in Fig. 7c, f, g and j.

Conducting a similar analysis for *κ* = 2, *σ*_*p*_ = 2 and *σ*_*h*_ = 0, as shown in the phase space in Fig. 4b, leads to similar observations emphasized above for the case of *κ* = 1 (see supplementary information, Fig. S1).

## V. CONCLUSION

Our main motivation in developing this work has been the paramount importance of the interactions between Tau proteins and microtubules in axons. In particular, Tau molecules play a crucial role in many neurodegenerative diseases referred to as *tauopathies*. Our goal was to study and describe how a stabilized microtubule can be decorated by a population of Tau in terms of coverage and spatial distributions. Based on published experimental evidences, we developed a model of Tau-microtubule interaction in which Tau proteins can bind to the microtubule lattice either along a protofilament (“*p*” mode) on two *αβ*-tubulin dimers or laterally (“*h*” mode) on two adjacent dimers as shown in Fig. 3. Within this framework, we show that line portraits in the phase space, {*ρ*_*p*_, *ρ*_*h*_}, of the microtubule decoration (see Fig. 4) are parameterized by the stochiometries of Taus with microtubule, *σ*_*p*_ and *σ*_*h*_, and defined by the ratio, *κ* = *k*_eq,p_*/k*_eq,h_, of equilibrium constants in the “*p*” and “*h*” modes, and that the location of points along those lines is controlled by the Tau:tubulin-dimers ratio, *x*, and the effective equilibrium constant, *k*_eff_ = *k*_eq,p_ + *k*_eq,h_. Each point in the phase diagram corresponds to a distribution of Taus attached on the microtubule wall which is characterized by two densities *ρ*_*p*_ and *ρ*_*h*_ and by an averaged distribution for the longitudinal spacing of Tau proteins, *P*_‖_(*r*). A microtubule decorated with Taus bound in a single mode (“*p*” or “*h*”) exhibits a single exponentional decay for *P*_‖_(*r*) (see Fig. 6c,d,g and h) while for the mixed case of Taus bound in two single modes (“*p*” and “*h*”) *P*_‖_(*r*) exhibits a multi-exponentional behavior (see Fig. 6k,l and Fig. 7d,e,h,i). From an experimental point of view, this model could be used as a theoretical framework to interpret and analyze binding data from cosedimentation assays and distributions for the longitudinal spacing of tau using quick-frozen, deep-etched suspension of microtubules as in [38], for example.

This work can be extended in several directions. A natural extension of this work would be to investigate the biological functions related to “*p*” and “*h*” binding modes. Indeed, taking into account that the binding domain of Tau involves three or four repeats [45] that are able to bind independently to a *α* or *β* monomer [27] and based on our model, one can speculate that Tau would adopt mostly an elongated form when bound along a protofilament and a more squashed form when bound across protofilaments. Therefore, by invoking arguments on the conformation adopted by Tau, the longitudinal and lateral form may have distinct biological functions as suggested by ref [37]. Moreover, due to the highly dynamic nature of Tau proteins even when bound to microtubules, the attachment geometry might be more complex than pure longitudinal or lateral modes. Indeed, in this work, we visualized a Tau-molecule as a stem of a given length and zero lateral extension but situations with Tau molecules having a nonzero extension or more complex shapes could also be investigated. To pursue this direction, the impact of the curvature of the microtubule on the decoration could also be studied. We expect that the helical geometry of the microtubule will affect mainly the Tau’s which are laterally bound, thus changing the ratio *κ* between the two equilibrium constants.

As an additional issue, it may turn out to be relevant to generalize the model and the approach outlined above to include a dynamical lattice, in order to study the effect of Tau on the microtubule dynamic instability. In the same spirit, the effect of a heterogeneous microtubule lattice with GTP and GDP states (heterogeneous binding sites) would need to be investigated as well.

## VI. METHODS

### Binding parameter estimates: *k*_eff_ and *x*

In human axons, the total concentration of Tau, [Tau], was found between ∼ 1% and 20% of the total tubulin-dimer concentration (both free and polymerized) [46]. In addition, more than 80% of the tubulins in the squid giant axon was found in the free form (i.e., not polymerized) [47]. In this specific case, the total concentration of tubulin is 5 times greater than the polymerized one i.e., [Tub_tot_] = 5 [Tub_poly_]. Throughout this work, we chose to work with conditions, 5 ≤ [Tub_tot_]/[Tub_poly_] ≤ 50 corresponding to a Tau:tubulin-dimer ratio, 0.1 ≤ *x* = [Tau]/[Tub_poly_] ≤ 10. The effective equilibrium constant can be estimated using the relation *k*_eff_ = [Tub_poly_]*/K*_*d*_ ≡ [Tau]/(*x* × *K*_*d*_) where *K*_*d*_ is the dissociation constant. Reported *K*_*d*_ values vary by more than two orders of magnitude from ∼ 0.01 *µ*M to ∼ 1 *µ*M [27, 28, 34, 41, 42, 48, 48–52]. Therefore, with a typical concentration of ∼ 1 − 2 *µ*M for Tau in axons [53, 54] and with the estimated ranges for *x* and *K*_*d*_, we end up with the range, 0.1 ≤ *k*_eff_ ≤ 10^3^.

### General decoration model: mathematical aspects

Expressions for the nearest neighbour distributions *P*_*k*=‖,⊥_(*r*) and insertion probabilities Φ_p,h_ have been derived by extending the approach based on the gap distribution as developed by McGhee and von Hippel [55] for an homogeneous infinite one-dimensional lattice. In what follows, assumptions underlying the method and derivation of *P*_*k*=‖,⊥_(*r*) and Φ_p,h_ will be presented.

#### Main approximation

The main idea for rendering tractable the calculations is to replace the two-dimensional microtubule lattice shown in Fig. 8 by two coupled linear lattices: protofilament and helix. The underlying approximation considers each direction as independent linear lattices. As illustrated in Fig. 8, for a protofilament “*j*” (helix “*i*”), Tau’s bound in “*p*” (“*h*”) mode cover (1 + *σ*_*p*_) [(1 + *σ*_*h*_)] consecutive sites while those bound in “*h*” (“*p*”) mode cover 1 site. According to this, the directional densities *ρ* _,*p*_ and *ρ* _,*h*_ can be expressed as a function of the microtubule coverages *ρ*_*p*_ and *ρ*_*h*_ as,

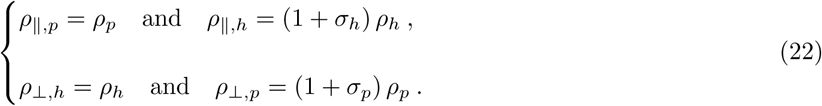

**FIG. 8:**
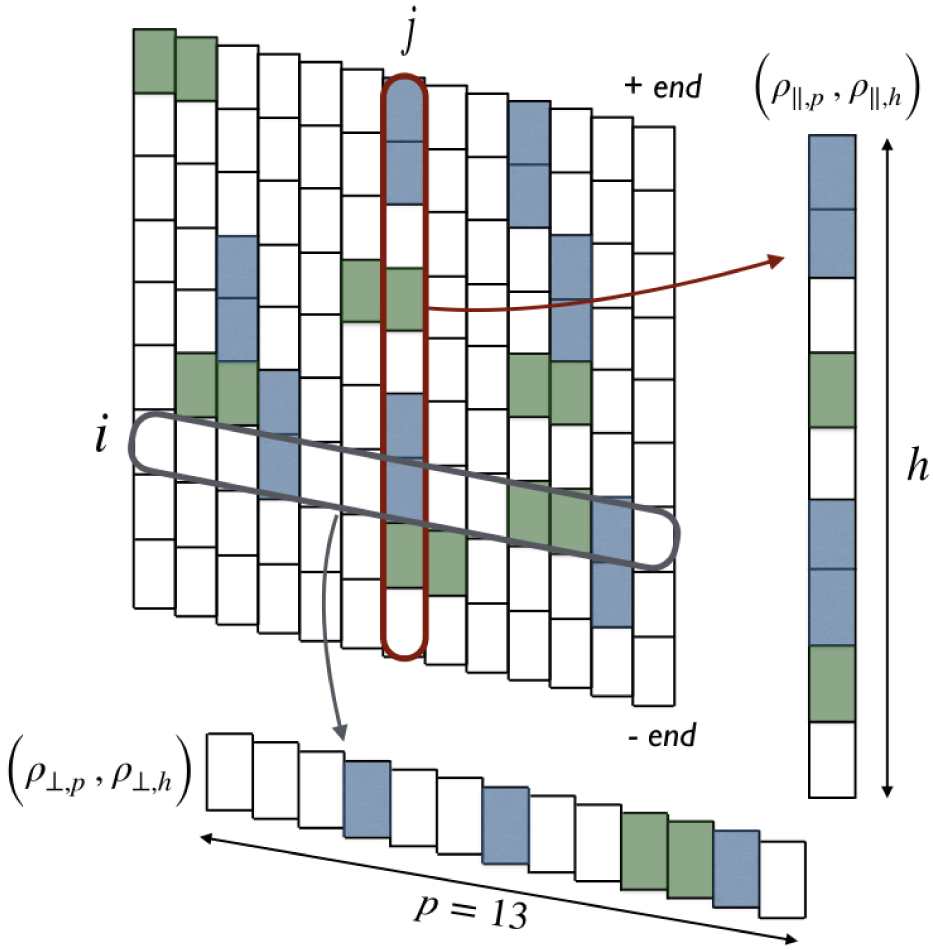
Illustration of the approximation consisting of replacing the two-dimensional lattice by two linear-lattices: (i) a helix *i* consisting of *p* = 13 sites with particles bound in “*h*” mode covering (1 + *σ*_*h*_) sites (in green) (density, *ρ*_⊥,*h*_) and particles bound in “*p*” mode covering 1 site (in blue) (density *ρ*_⊥,*p*_) and (ii) a protofilament *j* consisting of *h* sites with particles bound in “*p*” mode covering (1 + *σ*_*p*_) sites (in blue) (density *ρ* _‖,*p*_) and particles bound in “*h*” mode (in green) covering 1 site (density *ρ* _‖,*h*_).

#### Gap distribution

The gap distribution *f*_*k*=‖,⊥_(*g*) is defined as the probability of finding a gap consisting of *g* consecutive free lattice sites along a given direction *k*. As a consequence of the approximation shown in Fig. 8, we have to consider two gap distributions: *f*_‖_(*g*) for the protofilament linear lattice and *f*_⊥_(*g*) for the helicoidal linear lattice shown in Fig. 8 in red and blue, respectively. McGhee and von Hippel [55] have shown that *f*_*k*_(*g*) ∝ *u*_*k*_^*g*^ where *u*_*k*_ is the conditional probability that a lattice site selected at random along the direction *k* =‖,⊥ is free and that its adjacent site (the immediate top and right site for *k* =‖ and *k* =⊥, respectively) is free as well. Using the normalization condition of the probability for *f*_*k*_(*g*), we obtained:

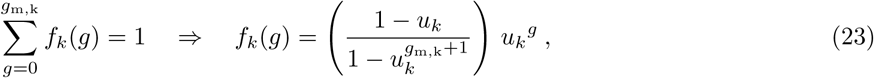

where *g*_m,k_ is the maximum physical gap between two bound Tau’s along the direction *k* = {‖, ⊥} given by,

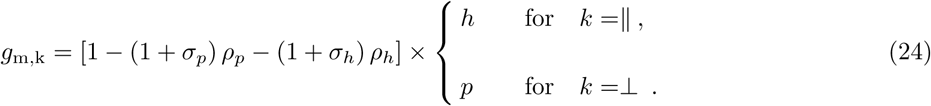

Every microscopic configuration being equally likely, the conditional probability *u*_*k*_ can be obtained as the ratio between the number of free sites, #(free sites)_*k*_ − 1, and the total number of objects (free sites plus bound Tau’s), #(free sites)_*k*_ + #(tau)_*k*_ − 1, forming the lattice along the direction *k* as,

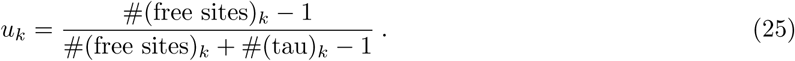

The −1 term in Eq.(25) accounts for that *u*_*k*_ is zero for a single site. Let *ℓ*_*k*_ (with *ℓ*_*k*_ = *h* and *ℓ*_*k*_ = *p* for *k* =‖ and *k* =⊥, respectively) be the total number of lattice sites along the *k* direction and, denote by *ρ*_free,k_ and *ρ*_*k*_ the densities of free lattice sites and bound Tau’s, respectively, along the direction *k*, given by,

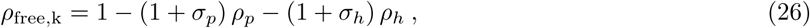

and

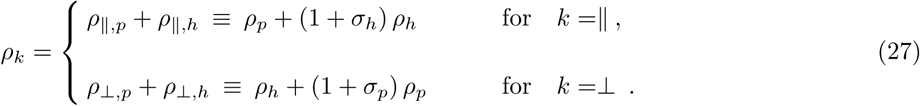

It follows that, #(free sites)_*k*_ = *ℓ*_*k*_*ρ*_free,k_ and #(tau)_*k*_ = *ℓ*_*k*_*ρ*_*k*_, such that,

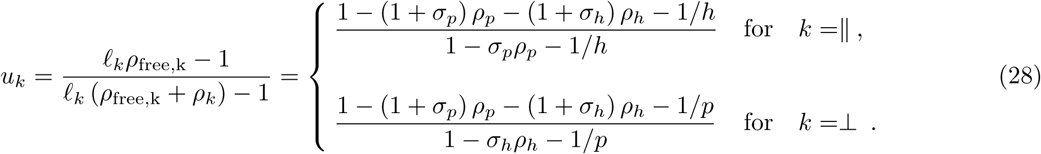

In the *h* ≫ 1 and *p* ≫ 1 limit, the gap distribution reads:

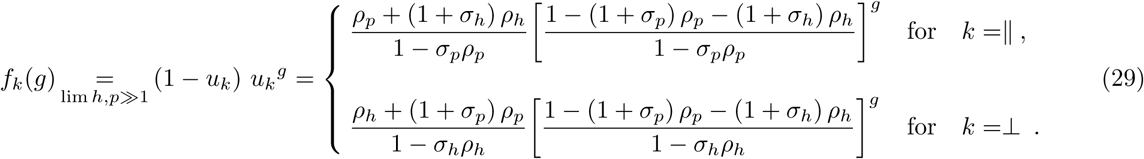

#### Nearest neighbour distribution and its first moment

The partial nearest neighbor distribution between Tau-molecules along the direction *k* =‖,⊥ is expressed as a function of the gap distribution as follows,

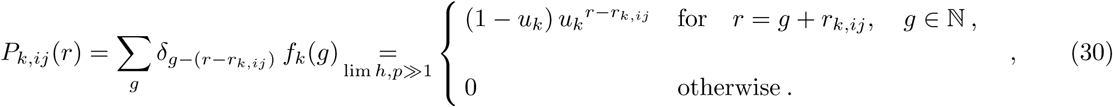

where the matrix elements *r*_*k,ij*_, correspond to the close packing center-to-center distance between Tau’s bound in “*i*” and “*j*” modes along the direction *k*, are given by

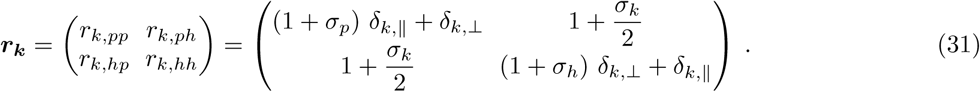

Using Eqs.(29) and (30) for *k* =‖ leads to *P*_‖,*ij*_(*r*) in Eq.(16) and for *k* =⊥, we have:

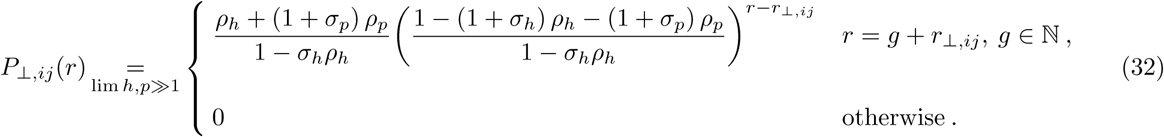

In the *h, p* ≫ 1 limit, the mean distance between two Tau-molecules along a *k* direction is obtained as,

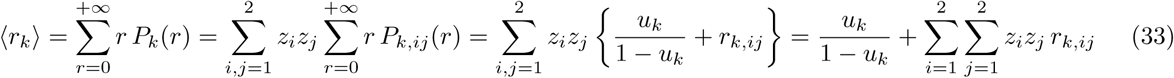

Finally, using the limit *h* ≫ 1 and *p* ≫ 1 in Eq. (28) leads to

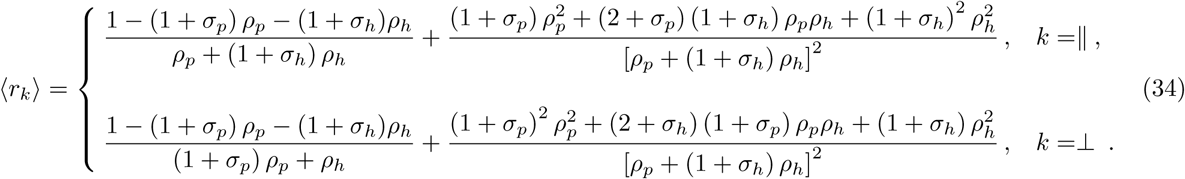

#### Insertion probabilities

The insertion probabilities Φ_p_ and Φ_h_ are obtained from, Φ_*k*_ = *n*_add,k_*/ℓ*_*k*_, where *ℓ*_*k*_ is the total number of lattice sites along the *k* direction (*ℓ*_*k*_ = *h* and *ℓ*_*k*_ = *p* for *k* =‖ and *k* =⊥, respectively) and *n*_add,*k*_ is the mean number of distinct ways for adding a Tau-molecule of size *σ*_*k*_ along the *k* direction (*σ*_*k*_ = *σ*_*p*_ and *σ*_*k*_ = *σ*_*h*_ for *k* =‖ and *k* =⊥, respectively). For a give number *n*_*k*_ of Taus bound along the *k* direction corresponds to a total of *n*_gap,*k*_ = *n*_*k*_ + 1 gaps, each of size *g* with a probability given by the gap distribution *f*_*k*_(*g*). Therefore, *n*_add,*k*_ reads as,

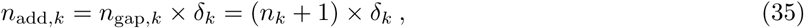

where *δ*_*k*_ counts the fraction of configurations allowing to accommodate a particle of size *σ*_*k*_ within each gaps. Following the approach in [55], *δ*_*k*_ is given by,

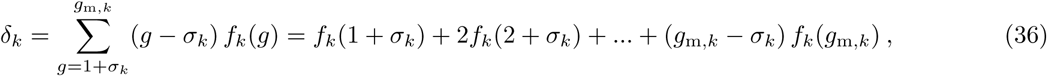

Eq.(36) indicates that for a particle of size *σ*_*k*_, there is one way of inserting that particle into a gap of size 1 + *σ*_*k*_ (with probability, *f*_*k*_(1 + *σ*_*k*_)), two ways into a gap of size 2 + *σ*_*k*_ (with probability, *f*_*k*_(2 + *σ*_*k*_)),, and so on up to the maximum physical gap size *g*_m,*k*_. The insertion probability is therefore given by,

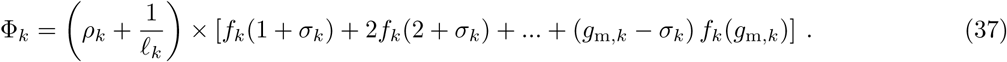

In the limit of a very long lattice (i.e., *ℓ*_*k*_ → ∞), Φ_*k*_ in Eq.(37) simply reduces to,

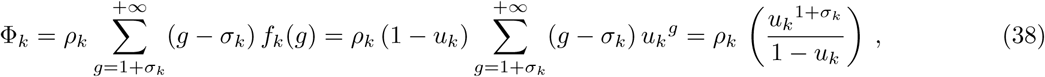

where *ρ*_*k*_ is given by Eq.(27) under the approximation described in VI. The Φ_p_ and Φ_h_ in Eq.(7) have been obtained by using Eq.(28) in the limit *h* ≫ 1 and *p* ≫ 1. A graphical representation of Φ_p_ and Φ_h_ for *σ*_*p*_ = *σ*_*h*_ = 1 are shown in Fig. 9.

**FIG. 9:**
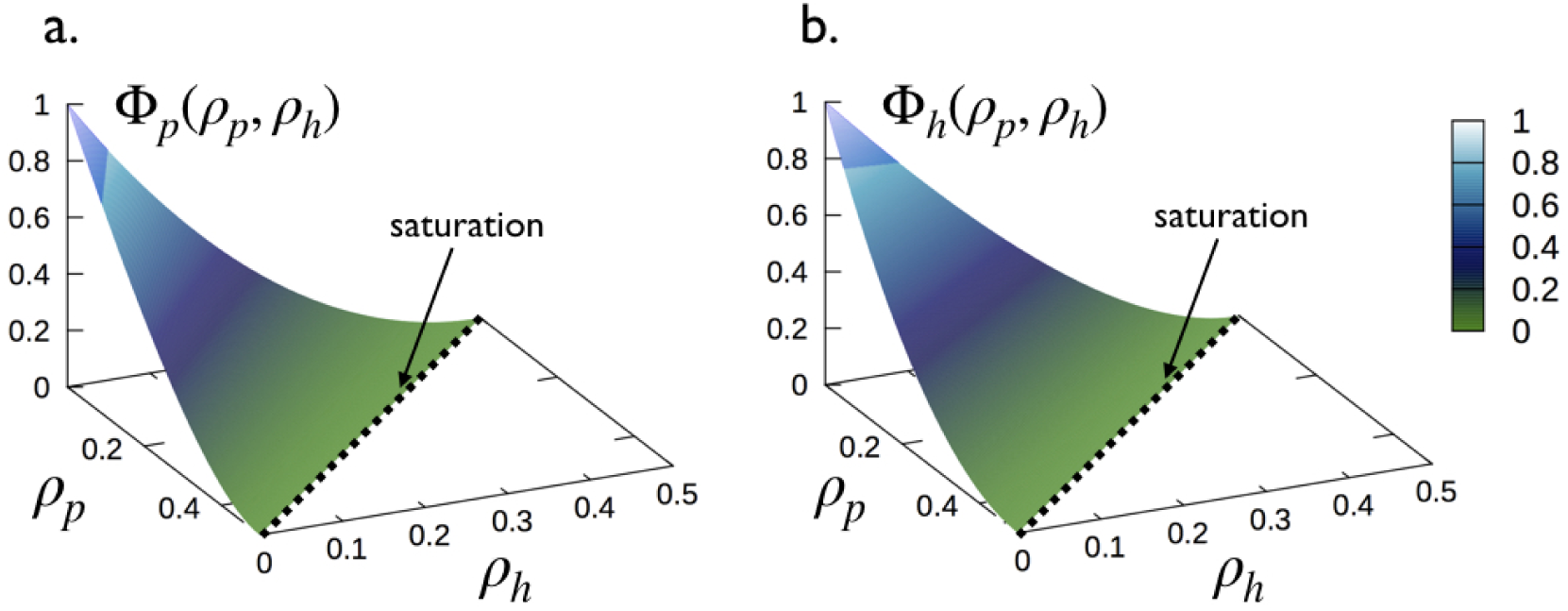
Surface plots of Φ_p_ and Φ_h_ in Eq.(7) as a function of *ρ*_*p*_ and *ρ*_*h*_ for *σ*_*p*_ = *σ*_*h*_ = 1. Dashed lines represent the saturation given by Eq.(8).

### Numerical solutions

The coupled system governing the evolution of densities *ρ*_*p*_ and *ρ*_*h*_ for the two populations of tau-molecules is obtained by using Eqs.(3) and (7) which lead to,

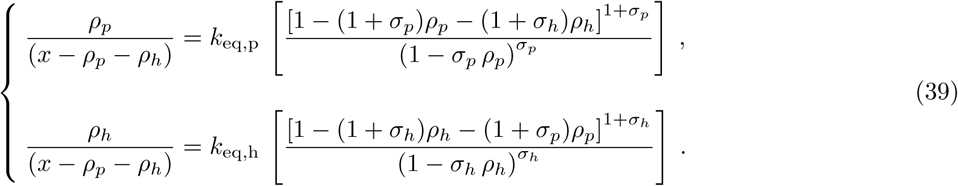

For a given set of parameters *σ*_*p*_, *σ*_*h*_, *k*_eq,p_ and *k*_eq,h_, Eq.(39) is solved for *ρ*_*p*_ and *ρ*_*h*_ using the function *fsolve* in MATLAB for several logarithmically spaced values of *x*. As shown in Fig. 10, each term in Eq.(39) corresponds to surfaces in the space (*ρ*_*p*_, *ρ*_*h*_) and their intersections leads to an unique point solution of Eq.(39). Numerical results for the case of interest *σ*_*p*_ = *σ*_*h*_ = 1 are discussed in Sec. IV.

**FIG. 10:**
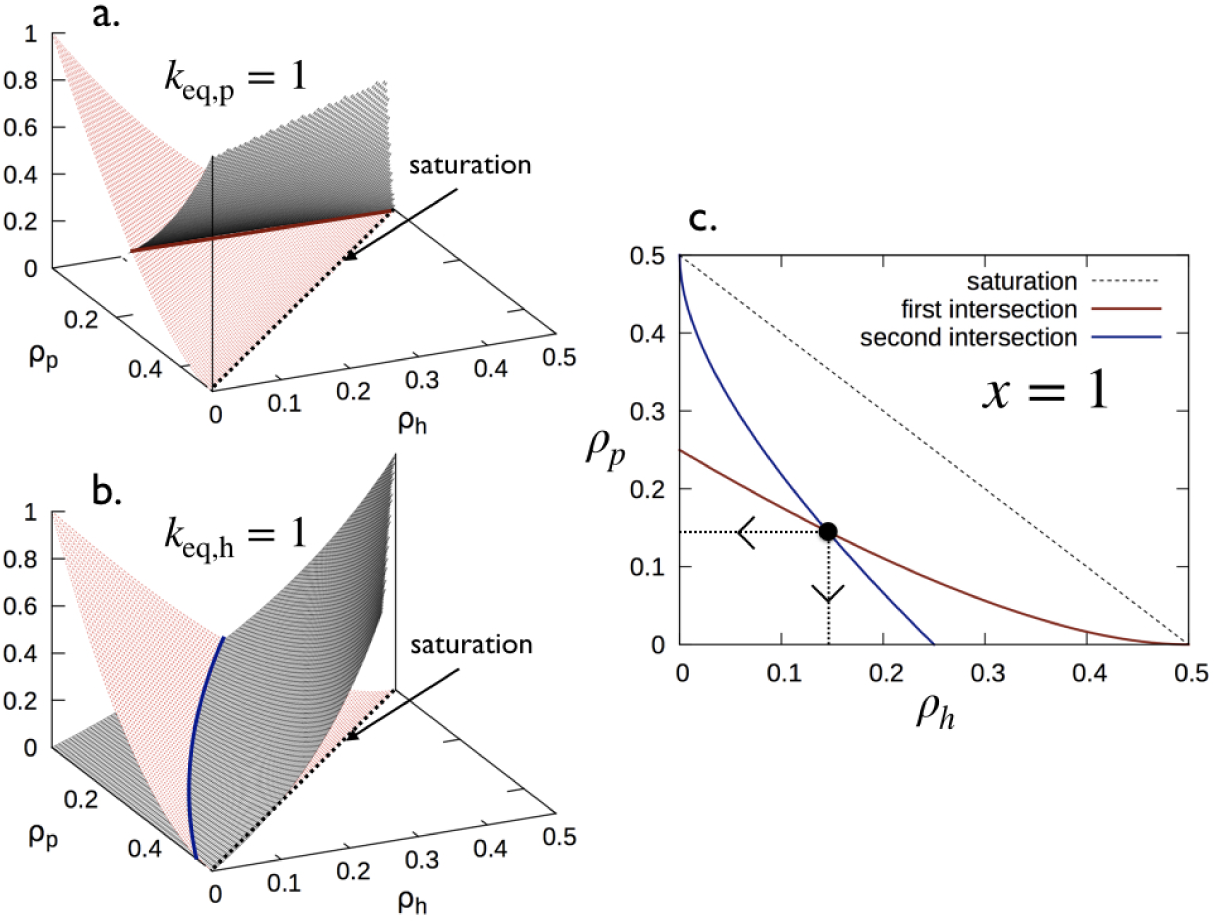
Graphical representation of the resolution of Eq.(39) for *ρ*_*p*_ and *ρ*_*h*_ for *σ*_*p*_ = *σ*_*h*_ = 1 with *x* = 1 and *k*_eq,p_ = *k*_eq,h_ = 1. (**a**,**b**) Surface plots corresponding to the first and second lines of Eq.(39). Dashed lines represent the saturation line given by Eq.(8). (**c**) Intersection between the two lines from Eq.(39) in (**a**,**b**) leading to an unique point solution of Eq.(39).

### Monte Carlo simulations

Stochastic simulations of the binding process described in III were performed for a two dimensional lattice of 615 × 13 ≈ 8000 sites corresponding to a 13−protofilament microtubule of about 615 × 8 *nm* = 4.92 *µm* long. Each data point of the coverage *ρ* shown in Fig. 6a has been obtained by averaging over 10^5^ simulated configurations. Histograms shown in Figs. 6c, d, g, h, k, and l and 7d, e, h, and i have been obtained by computing all the center-to-center distances between nearest neighbors along the protofilament direction (*k* =‖) averaged over 10^5^ simulated configurations while the snapshots in Fig. 6b, e, f, i, j and m and in Fig. 7c, f, g and j show a zoom in (13 × 40 sites) of a single simulated configuration.

## Acknowledgments

We thank Dr. Timothy Ziman for revising language across whole manuscript. J.H is a PhD student supported by a grant from the Ministry of Education and Research of France through the Ecole Doctorale de Physique de Grenoble (ED n°47) of Grenoble Alpes University.

## Data Availability

The datasets generated and analysed during the current study are available from the corresponding author on reasonable request.

## Author Contributions

J.H and D.B contributed equally to this work.

## Additional Information

**Supplementary information** accompanies this paper at:

## S1. BIBLIOGRAPHIC DATA BASE

### A. Interaction tau-microtubule

The list of 54 papers below related to the interaction tau-microtubule have been obtained with <monospace>PubMed</monospace> and <monospace>GoogleScholar</monospace> by using the following list of keywords: (i) *Tau microtubule binding site*, (ii) *Tau microtubule interaction*, (iii) *Tau microtubule topology*, (iv) *Tau microtubule binding modes*, (v) *Tau microtubule binding parameters*, (vi) *Tau microtubule binding stoichiometry*, (vii) *Tau microtubule binding dissociation constant*, (viii) *Tau microtubule assembly domain*, (ix) *Tau microtubule decoration*, (x) *Tau binding domain*, (xi) *Tau microtubule functions* and (xii) *Tau tubulin complex*.

### B. Structure of the microtubule lattice

The list of 16 papers below related to the structure of the microtubule lattice have obtained with `PubMed` and `GoogleScholar` by using the following list of keywords: (i) *Microtubule structure*, (ii) *Microtubule lattice*, (iii) *Microtubule surface lattice* and (iv) *Microtubule lattice seam*.

## S2. MICROTUBULE COVERAGE FOR *σ*_*p*_ ≠ *σ*_*h*_

**FIG. S1:**
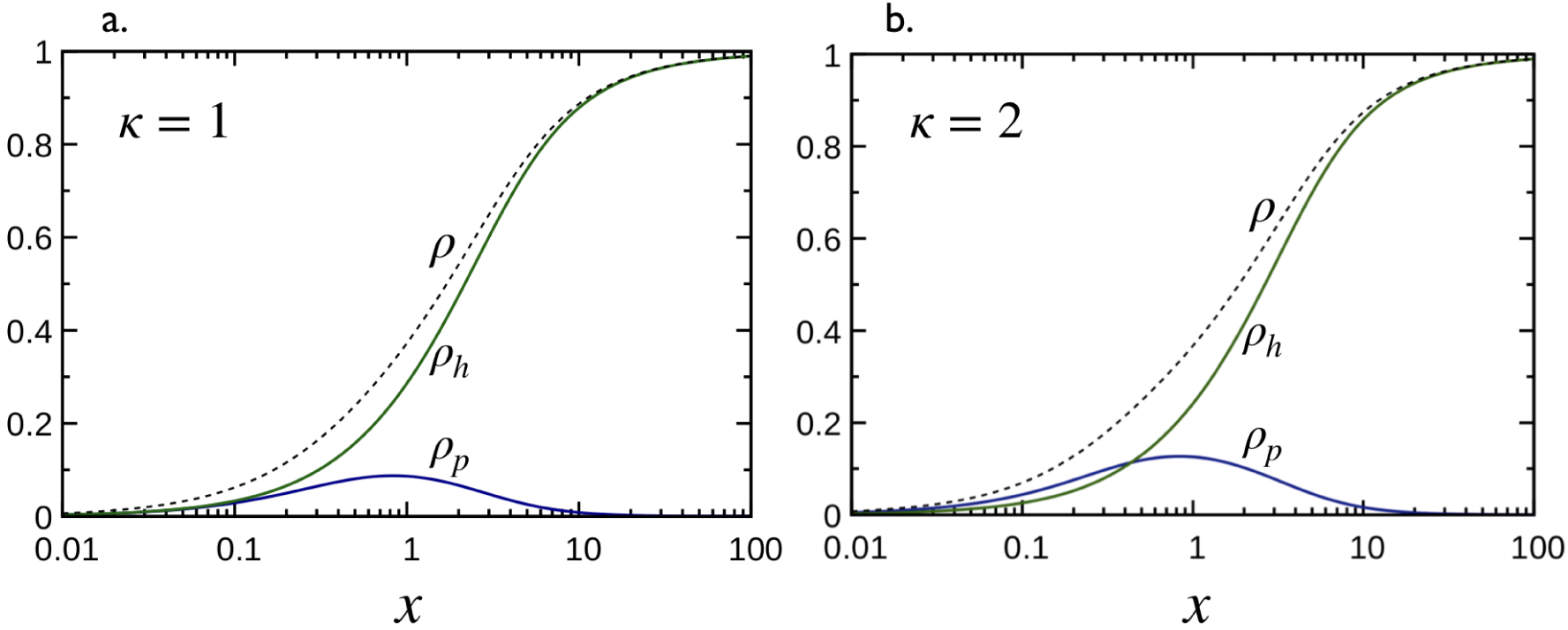
Decoration of a microtubule by Tau proteins for *σ*_*p*_ = 2 and *σ*_*h*_ = 0. Densities *ρ*_*p*_, *ρ*_*h*_ (solid lines) and the total coverage *ρ* = *ρ*_*p*_ + *ρ*_*h*_ (dasheh lines) as a function of *x* for *k*_eq,p_ = *k*_eq,h_ = 1 (i.e., *κ* = 1) in (**a**) and *k*_eq,p_ = 2, *k*_eq,h_ = 1 (i.e., *κ* = 2) in (**b**). Solid lines are obtained from the numerical solution of Eq.(3) (see manuscript).

## Footnotes

[*] Eq.(8) is an approximation to the saturation for the binding in “*h*” mode because the binding at the MT seam is not allowed. Therefore, for *σ*_*h*_ = 1 and 13-protofilaments MT, the highest density in “*h*” mode is *ρ*_*h*_ = 6/13 ≈ 0.46, instead of 1/2, corresponding to 6 Taus bound in “*h*” mode (i.e., 12 lattice binding sites are occupied) per MT helix. The correct saturation line is shown in Fig. 4a in dashed line and is given by: *ρ*_*h,s*_(*ρ*_*p,s*_) = 6/13 for 0 ≤ *ρ*_*p,s*_ *<* 1/26, and, *ρ*_*h,s*_(*ρ*_*p,s*_) = 1/2 − *ρ*_*p,s*_ for 1/26 ≤ *ρp,s* ≤ 1/2 As it can be seen in Fig. 4, the saturation line in Eq.(8) is exact for 1/26 ≤ *ρp,s* ≤ 1/2 but becomes an approximation for *ρp,s <* 1/26 yielding a maximum relative error of about 4%.

